# Striatal dopamine synthesis capacity in Parkinson’s disease: Effects of age, sex, and body mass index in a large [^18^F]fluorodopa PET cohort

**DOI:** 10.1101/2025.10.13.681966

**Authors:** Tuulia Malén, Jouni Tuisku, Marco Bucci, Severi Santavirta, Valtteri Kaasinen, Sakari Kaasalainen, Janne Isojärvi, Jarmo Hietala, Juha Rinne, Lauri Nummenmaa

## Abstract

**BACKGROUND:** In Parkinson’s disease (PD), the degeneration of nigrostriatal dopaminergic neurons leads to motor symptoms. Positron emission tomography (PET) using radioligand [^18^F]fluorodopa detects reduced striatal dopamine synthesis capacity in PD patients. Demographic factors such as sex and BMI are also associated with dopamine synthesis capacity. The combined contribution of demographic and clinical effects however remains elusive. It also remains unresolved how the dopamine synthesis capacity is correlated between striatal subregions and how the dopamine synthesis capacity and dopamine receptor function across striatal regions are associated with each other in PD patients and healthy controls.

**MATERIAL, AIMS, AND METHODS:** For this retrospective register-based study, we used baseline [^18^F]fluorodopa PET data acquired at the Turku PET Centre between the years 1988-2016 with three different scanners (Ecat 931, GE Advance, HRRT). The data included scans of 350 adult human subjects, including 132 healthy controls (65 males and 67 females), and 218 PD patients (134 males and 84 females). The primary aim was to simultaneously investigate the effects of PD, age, sex and BMI on regional dopamine synthesis capacity (influx rate constant Ki^ref^ quantified with Patlak) using Bayesian linear regression. Secondary aims were to assess interregional correlations of dopamine synthesis capacity, and the association between regional presynaptic dopamine synthesis and postsynaptic dopamine type 2 receptor (D_2_R) availability in subjects who also had a proximal [^11^C]raclopride PET scan.

**RESULTS:** We found modest support for age-dependent decline in dopamine synthesis capacity, increased capacity for higher BMIs, and for higher capacity in females versus males. These effects were smaller than the effect of PD status. Dopamine synthesis capacity across regions was correlated in both patients and controls. Support for positive correlation between the synthesis capacity and the D_2_R was observed in caudate nucleus.

**CONCLUSIONS:** PD and demographic effects are independently associated with the striatal dopamine synthesis capacity. The capacity is reduced by PD, decreased through age (particularly after the age of around 50), higher in females versus males, and weakly increased in higher BMIs. Synthesis capacity is correlated between the striatal and thalamic regions in both PD patients and controls. The dopamine synthesis capacity was positively correlated with the D_2_R availability in caudate. Scanner affects the estimates of dopamine synthesis capacity measured with [^18^F]fluorodopa PET, and it is preferrable to adjust for such variation in the data.

## Introduction

Parkinson’s disease (PD) is characterized by progressive degeneration of nigrostriatal dopaminergic neurons, leading to initially asymmetric motor symptoms and reduced striatal dopamine synthesis capacity detectable with [^18^F]fluorodopa positron emission tomography (PET) (Kalia & Lang, 2015). [^18^F]fluorodopa uptake is a measure of aromatic L-amino acid decarboxylase (AADC) activity reflecting dopamine synthesis capacity, and it reliably distinguishes patients with PD from healthy controls even at early disease stages (Kaasinen & Vahlberg, 2017).

Normal ageing is also associated with gradual dopaminergic decline (Malén et al., 2022), consistently demonstrated for dopamine receptors and transporters, although evidence for changes in dopamine synthesis capacity is less consistent (Karrer et al., 2017). Some studies suggest sex differences, with females showing higher [^18^F]fluorodopa uptake than males, and potential age-related reductions particularly in men (Laakso et al., 2002). Moreover, body mass index (BMI) has been linked to striatal dopamine function, with higher BMI associated with lower dopamine synthesis capacity (Janssen & Horstmann, 2022) while dopamine receptor availability is unaffected (Pak & Nummenmaa, 2023).

Previous [^18^F]fluorodopa PET studies have been limited by modest sample sizes and restricted scope, focusing either on PD versus controls or on demographic effects in healthy subjects. The combined contribution of demographic influences and PD-related changes has not systematically been addressed. Furthermore, the extent to which dopamine synthesis capacity is coupled within striatal subregions and dopamine receptor function across striatal regions in PD and controls remains unknown. Earlier evidence from healthy controls suggests a positive association between striatal dopamine synthesis capacity and type 2 dopamine receptor (D_2_R) availability (Berry et al., 2018).

The aim of this study was to pool retrospective [^18^F]fluorodopa data from a large (350 subjects) sample of subjects to simultaneously assess the effects of PD, age, sex and BMI on striatal dopamine synthesis capacity. Secondary aims were to investigate: (1) interregional correlations of dopamine synthesis capacity, (2) the association between presynaptic dopamine synthesis and postsynaptic D_2_R availability in subjects with dual-tracer scans, and (3) methodological factors including scanner effects and atlas-versus MRI-based quantification approaches. Our final aim was to provide the mean dopamine synthesis brain maps of healthy controls and PD patients in NeuroVault (https://neurovault.org).

## Material and methods

### Data

The data were baseline [^18^F]fluorodopa PET images of 350 adult human subjects, including 132 healthy controls (65 males and 67 females), and 218 PD patients (134 males and 84 females) for whom sufficient demographic and imaging data were available. The scans were acquired at the Turku PET Centre between the years 1988-2016 with three different scanners (Ecat 931, GE Advance, HRRT, from oldest to newest at the site). See details for the original studies in the supplementary material (section **Publications whose data are used in the current study**). The imaging and the demographic data were derived from Aivo-database (https://aivo.utu.fi) maintained at the site.

The initial sample of 367 subjects was cut down to 350 after exclusion criteria were applied to 17 subjects as follows: one PD patient with prior thalamotomy, one with clinically undetermined parkinsonian syndrome after follow-up, and one with later determined drug-induced parkinsonism. Additionally, we excluded seven subjects due to insufficient number of collected frames for reliable modelling (four or less within 15-60 minutes of the scanning), six with missing PET estimates from one or more regions of interest (ROI), and one with abnormally low signal intensity likely reflecting failed tracer injection and/or preprocessing error. After these exclusions, the total sample consisted of 350 subjects.

The age range of the subjects was 19-81 years. Body mass index (BMI) data were available for 220 subjects (63% of the total sample), ranging from 17.8 to 43.8 (BMI subsample). Information on smoking status, handedness, PD duration, motor symptom severity, and medication use was not systematically available. Among the healthy controls, 14 individuals in the total sample (11 in the BMI subsample) were non-psychotic first-degree relatives of patients with schizophrenia. Descriptive statistics of the total sample and BMI subsample are summarized in **Tables 1** and **2**, respectively. Age and BMI distributions are shown in **Figure 1**.

**Table 1.**
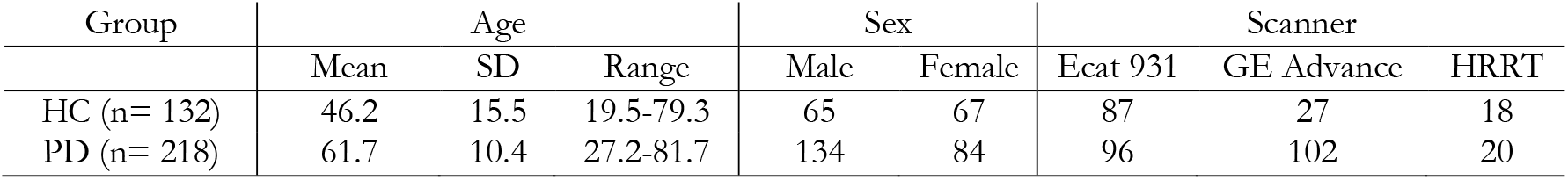
Demographic and imaging characteristics of the total sample (n= 350). HC= healthy controls, PD= patients with Parkinson’s disease, SD= standard deviation. Scanner column shows the number of subjects imaged with each scanner.

**Table 2.** Demographic and imaging characteristics of the subsample of subjects with BMI information available (BMI subsample, n= 220, 63% of the total sample). HC= healthy controls, PD= patients with Parkinson’s disease, SD= standard deviation. Scanner column shows the number of subjects imaged with each scanner.

| Group | Age (years) |  |  | Sex |  | BMI |  |  | Scanner |  |  |
| --- | --- | --- | --- | --- | --- | --- | --- | --- | --- | --- | --- |
|  | Mean | SD | Range | Male | Female | Mean | SD | Range | Ecat 931 | GE Advance | HRRT |
| HC (n= 94) | 45.5 | 15.3 | 19.5-79.3 | 46 | 48 | 25.3 | 3.5 | 17.8-38.8 | 52 | 26 | 16 |
| PD (n= 126) | 62.8 | 10.0 | 33.0-79.7 | 78 | 48 | 26.2 | 3.9 | 18.7-43.8 | 15 | 101 | 10 |

**Figure 1.**
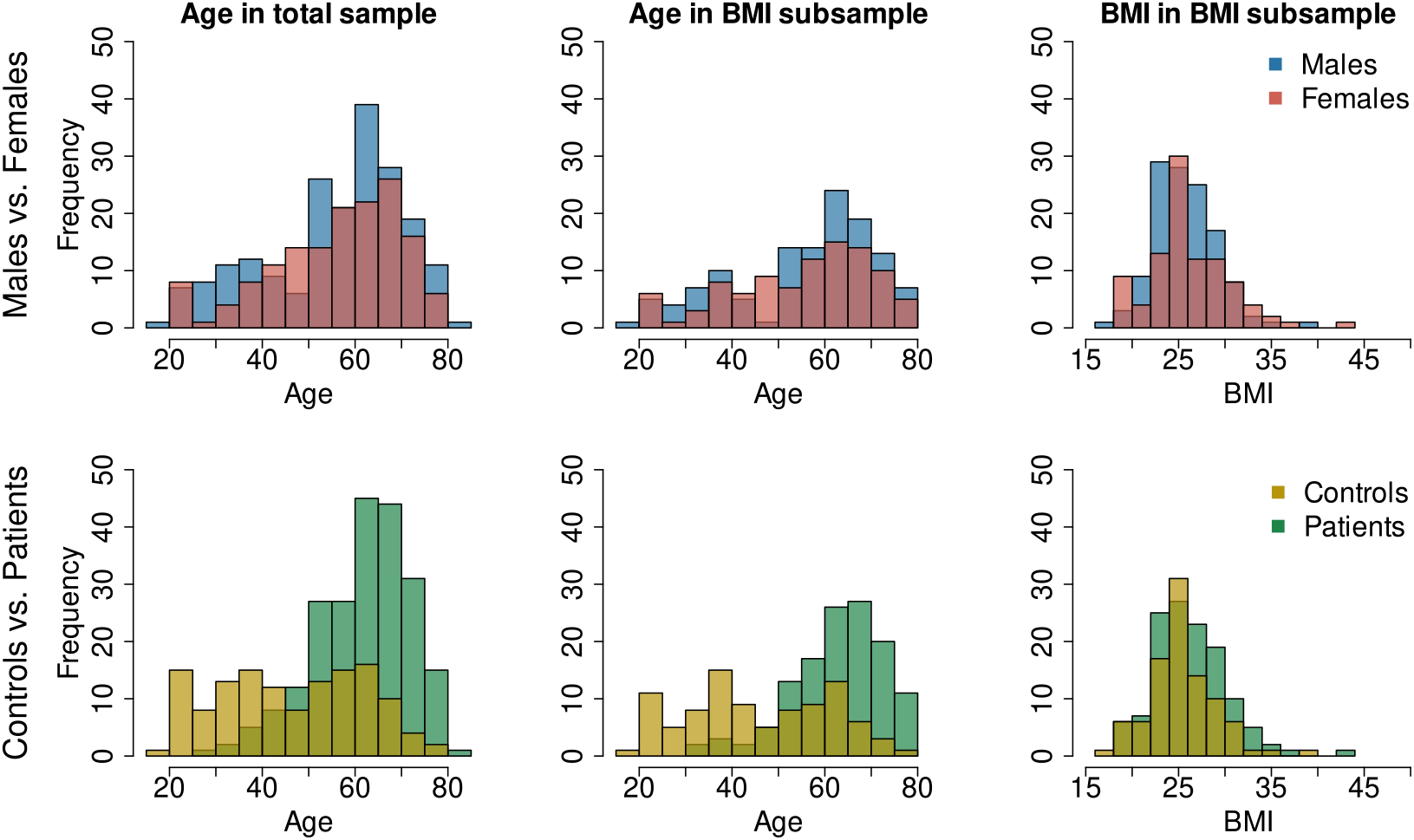
Age and BMI histograms of the total sample and the subsample of subjects with BMI information available (BMI subsample, n= 220, 63% of the total sample). The darker shades of red and yellow are overlapping with blue and green, respectively.

PET scan using [^11^C]raclopride and taken within ±48 days from the [^18^F]fluorodopa scan was available from 6 healthy controls and 26 PD patients. The subjects were included in the sample of our study on D_2_R (Malén et al., 2024). The scanner for all 32 [^18^F]fluorodopa and the matching 32 [^11^C]raclopride images was Ecat 931. Out of those for whom we had the information of the PD duration available (18/26), four subjects were in the early stage of PD (< 4 years from diagnosis) and the remaining 14 were in later stage (> 4 years from diagnosis) at [^18^F]fluorodopa imaging. Descriptive statistics of the subjects are given in **Table 3**.

**Table 3.** Demographic and clinical characteristics of the subsample of subjects with closely (maximum of 48 days apart) imaged [^18^F]fluorodopa and [^11^C]raclopride scans available (n= 32). HC= healthy controls, PD= patients with Parkinson’s disease, SD= standard deviation, NA= not applicable. Duration= Parkinson’s disease duration from diagnosis of 18 patients with the available information. The table shows information at [^18^F]fluorodopa scan. BMI is not given in the table, as it was available only for two subjects.

| Group | Age (years) | | | Sex | | Duration (years, $n = 18$) | | |
| --- | --- | --- | --- | --- | --- | --- | --- | --- |
|  | Mean | SD | Range | Male | Female | Mean | SD | Range |
| HC ($n = 6$) | 60.6 | 7.0 | 51.2-72.2 | 2 | 4 | NA | NA | NA |
| PD ($n = 26$) | 58.8 | 12.8 | 34.2-81.7 | 15 | 11 | 11.9 | 7.7 | 1-29 |

### PET image preprocessing

The [^18^F]fluorodopa uptake was quantified using the Patlak method where the influx rate constant Ki^ref^ (Gjedde et al., 1991) was estimated using multiple time graphical analysis (Patlak & Blasberg, 1985) in regional and voxel level within 15 to 60 minutes by using the occipital cortex as reference region (Nurmi et al., 2001). Ki^ref^ is a slope that reflects the efficiency of AADC activity, i.e. dopamine synthesis capacity (Kaasinen & Vahlberg, 2017). Especially in regions with particularly low Ki^ref^, the Patlak-based Ki^ref^ estimate may be below zero, possibly due to less signal with noisy filtered back projection reconstruction. In the total sample, there were altogether 29 negative Ki^ref^ estimates (4 in caudate, 1 in accumbens, 7 in putamen, and 17 in thalamus, involving observations of 20 subjects), of which 27/29 were acquired with Ecat 931 and 21/29 from PD patients. We did not remove observations with negative Ki^ref^ estimates to avoid a positive bias on the mean. The negative Ki^ref^ estimates could also stem from metabolites (Sossi et al., 2003). However, all subjects were administered 150 mg of carbidopa (a peripheral AADC inhibitor) an hour prior to tracer injection to minimize peripheral metabolism (Elsinga et al., 2006; Joutsa et al., 2015).

Image preprocessing and modelling were performed with an in-house automated image analysis pipeline Magia (Karjalainen et al., 2020) running on MATLAB (mathworks, Inc, MA, USA). The pipeline included PET frame-by-frame realignment and for those cases with available magnetic resonance imagen (MRI), co-registration of MRI and PET images. The rate of dopamine synthesis can be measured by quantifying the accumulation of [^18^F]fluorodopa in the striatum (Volkow et al., 1996). Data were analyzed in striatal regions of interest (ROIs; caudate nucleus later referred to as caudate, nucleus accumbens later referred to as accumbens, putamen), as well as thalamus. Bilateral ROIs were defined using Harvard-Oxford atlas, which was transformed into subject native space using an in-house created PET-template. For subjects with structural MRI data available (n=45), anatomical ROIs were extracted using FreeSurfer (version 7.2.0, https://surfer.nmr.mgh.harvard.edu/). For validation purposes, we compared the Ki^ref^ estimates in the subsample of subjects where both MRI-based and atlas-based normalization was possible. The one-tailed (greater) Pearson correlation between the Ki^ref^ estimates of the two alternative methods was 0.57 (Spearman 0.69) in caudate, 0.13 (Spearman 0.40) in accumbens, 0.79 (Spearman 0.79) in putamen, and 0.46 (Spearman 0.45) in thalamus. The Pearson and Spearman correlations were statistically significant (p-values <= 0.001), except in accumbens. More details of the validation in supplementary material section **Comparison of atlas-based and MRI-based Ki**^**ref**^ **estimates**. The uptake of [^11^C]raclopride was quantified as described in our previous D_2_R study (Malén et al., 2024).

### Statistical analysis

To manage, analyze and visualize our data, we used MATLAB (The Mathworks Inc., 2021), MRIcroGL (https://www.nitrc.org/projects/mricrogl), and R (R Core Team, 2025) in RStudio (Posit team, 2023), including the packages data.table (Dowle, 2023), tidyr (Wickham, 2021), dplyr (Wickham, 2023), stringr (H. Wickham, 2022), reshape2 (Wickham, 2007), scales (H. S. Wickham, Dana, 2022), ggplot2 (Wickham, 2016), gridExtra (Auguie, 2017), ggdist (Kay, 2023a), patchwork (Pedersen, 2024), and tidybayes (Kay, 2023b).

For statistical analyses, mean Ki^ref^ estimates across hemispheres were used, as atlas-based normalization may not adequately capture subject-specific asymmetry. To ensure that potential lateralization effects were not overlooked, hemispheric differences were assessed in the subset with MRI-based Ki^ref^ estimates derived separately for left and right ROIs. Results supporting the validity of mean hemispheric estimates are presented in the supplementary material (section **Lateralization of Ki**^**ref**^ **estimates**).

#### Modeling of age, sex, BMI, PD, and scanner effects

We conducted Bayesian linear regression modeling using the R package brms (Bürkner, 2017, 2018, 2021) utilizing RStan (Stan Development Team, 2023). We ran the models separately for each ROI (caudate, accumbens, putamen, and thalamus). In the regional models, we estimated the fixed, population-level main effects of standardized age (slope), sex (difference between males and females), standardized BMI (slope), and PD (difference between PD patients and healthy controls) on standardized Ki^ref^.

Despite calibration and standardization, different PET scanners used for each study may induce variation in the data, because the scanners differ in their properties, such as sensitivity and resolution, which may affect the acquired Ki^ref^ estimate. To address this potential variation, we also estimated the fixed (population-level) main effect of scanner on the standardized Ki^ref^, as we wanted to estimate the scanner effect, as well as the effects of age, sex, BMI, and PD independently of the scanner.

We wanted to maximize the number of subjects in the modeling. Thus, we primarily estimated the effects of age, sex, PD, and scanner in the total sample, but validated these effects also in the BMI subsample. The fixed effect of BMI was estimated only in the subsample where BMI was known.

In the Bayesian modeling, a prior probability distribution (prior) is given to each parameter in the model (McElreath, 2020). Each prior is updated with the data, resulting in a posterior distribution (posterior) that describes the probability distribution of the parameter (McElreath, 2020). In the regression modeling, we were interested in the posteriors for the regression coefficients of our fixed effects. Primarily, we set normal prior with expected value of 0, and standard deviation (SD) of 1 for the regression coefficients (for age, sex, BMI, clinical status of the subject as PD patient or healthy control, and scanner) and used the default priors of brms for the remaining parameters.

To assess the effect of the chosen priors, we ran separate regional models using alternative priors that were assigned to the regression coefficients of BMI, sex, and the clinical status of the subject as PD patient or healthy control. The alternative priors for BMI, sex and PD were normal distributions weighting negative association (expected value: -1, SD: 0.5). The alternative prior specification was based on previous findings of lowered dopamine synthesis capacity in high BMI (Janssen & Horstmann, 2022), in males versus females (Laakso et al., 2002), and in PD patients versus healthy controls (e.g. Cropley et al., 2008; Pavese et al., 2011; Samii et al., 1999). Using alternative priors did not substantially change the findings. Normality and homoscedasticity of the modeling residuals suggest that the assumptions of the linear regression model are met, and that the model fits the data sufficiently. Assessing the residuals, we did not observe clear violations of normality nor homoscedasticity. More details of the diagnostics (including the modeling with alternative priors and the assessment of residuals), are given in the supplementary material (section **Model diagnostics**).

In the primary modeling, we calculated the age effect on Ki^ref^ for males and females together. To assess possible sex-dependency in the age-effect, we estimated the effect of age for males and females independently (fixed interaction of age and sex) in separate regional models (see details in supplementary material section **Interaction of age and sex**).

In the primary regression modeling, we forced the associations between continuous predictors (age and BMI) and the dependent variable (Ki^ref^) as slopes (straight lines). To assess whether the linear function adequately approximates the age and BMI effects, we ran separate regional models allowing nonlinearity in these effects (see details in supplementary material section **Linearity assessment of the age and BMI effects**).

We specified each regression model to run with four chains (Markov chain Monte Carlo, MCMC), and each chain was defined with 4000 iterations including 1000 warm-ups. Otherwise, the default specifications of brms were applied, except for the models allowing nonlinear effects, adapt-delta and maximum treedepth were modified to facilitate model convergence. Using these model fitting specifications, the models showed neither divergent transitions nor Rhats > 1, supporting sufficient model convergence (Vehtari et al., 2021). The R syntax for running the regression models is presented in the supplementary material (**R syntax for the regression models**).

#### Interregional correlations of Ki^ref^

We analyzed interregional associations in Ki^ref^ separately in healthy controls and in PD patients in the total sample (n= 350). The Pearson correlations were analyzed separately for each scanner to avoid confounding effect of scanner. We tested whether the Ki^ref^ was normally distributed separately in healthy controls and PD patients within each scanner and each ROI using Shapiro-Wilk testing (significance level 0.01) in R package stats (R Core Team, 2025). Most of the tests (15/24) did not conflict with the null-hypothesis of normally distributed data, and we decided to estimate the correlation using (two-sided) Pearson correlation with default specification of cor.test function in R package stats (R Core Team, 2025). We used original scale and not standardized Ki^ref^ estimates, as standardization here does not facilitate the interpretation of the findings similarly as in the regression modeling.

#### The relationship between striatal dopamine synthesis and and D_2_R availability

For the 32 subjects with baseline [^18^F]fluorodopa and [^11^C]raclopride scans conducted less than 48 days apart, we calculated the Pearson correlation (two-tailed testing) between [^18^F]flurodopa Ki^ref^ and [^11^C]raclopride binding potential (BPND) estimates, to assess the baseline link between regional dopamine synthesis capacity and D_2_R receptor availability. Based on Shapiro-Wilk testing (significance level 0.01), the regional Ki^ref^ and BPND estimates did not conflict with the null-hypothesis of normally distributed data (except in thalamus). The relationship between Ki^ref^ and BPND was visually assessed not only for the subjects together but also separately for healthy controls and PD patients to give more insight on whether the relationship differs between the groups. In the assessment altogether, we used the original scale estimates.

## Results

### Group-differences in Ki^ref^

Regional Ki^ref^ estimates (**Figure 2**), and the group-specific whole-brain maps for mean Ki^ref^ (**Figure 3**) demonstrate differences between healthy controls and PD patients. The whole-brain coefficient of variation -maps show that the variation of Ki^ref^ estimates is greatest in thalamus (**Figure 3**). Scanner-specific distributions are shown in **Figure 4**.

**Figure 2.**
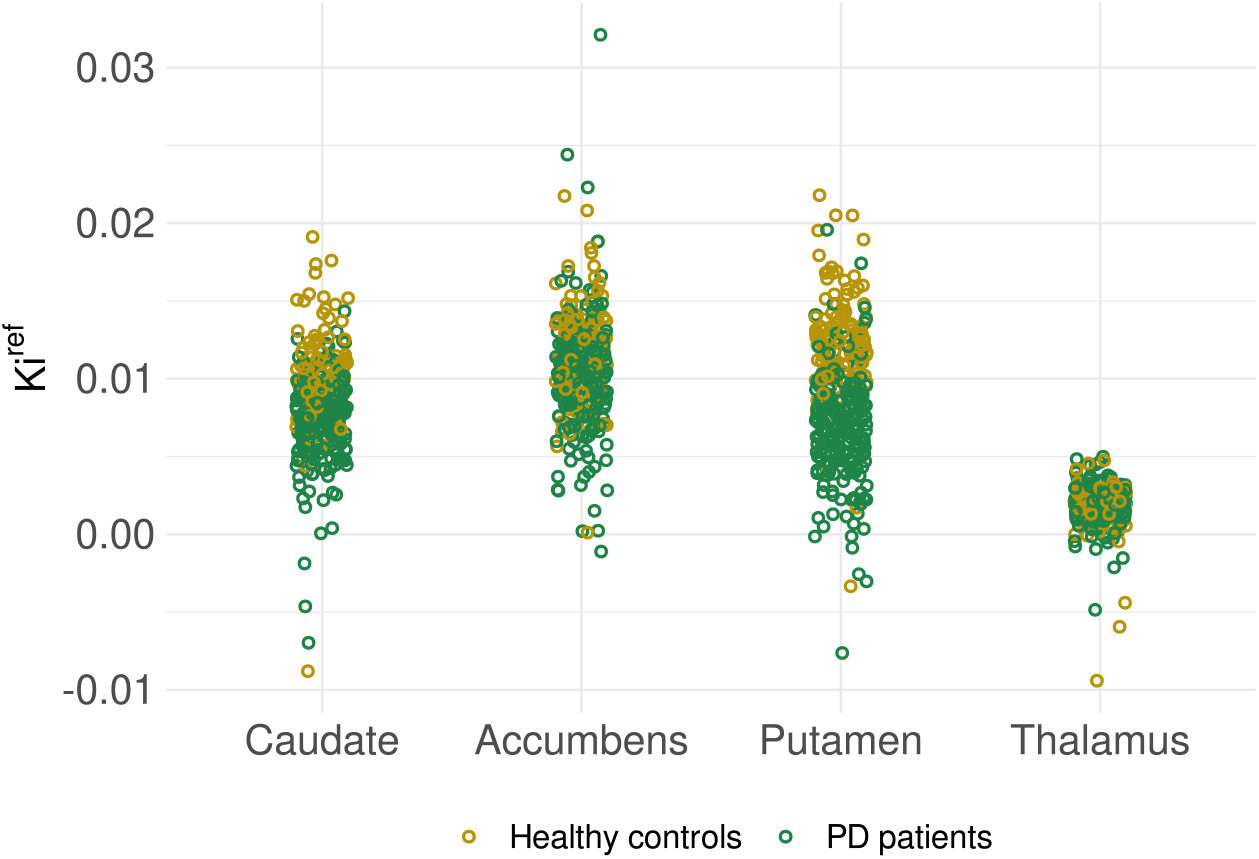
Regional Ki^ref^ estimates in healthy controls and PD patients in the total sample.

**Figure 3.**
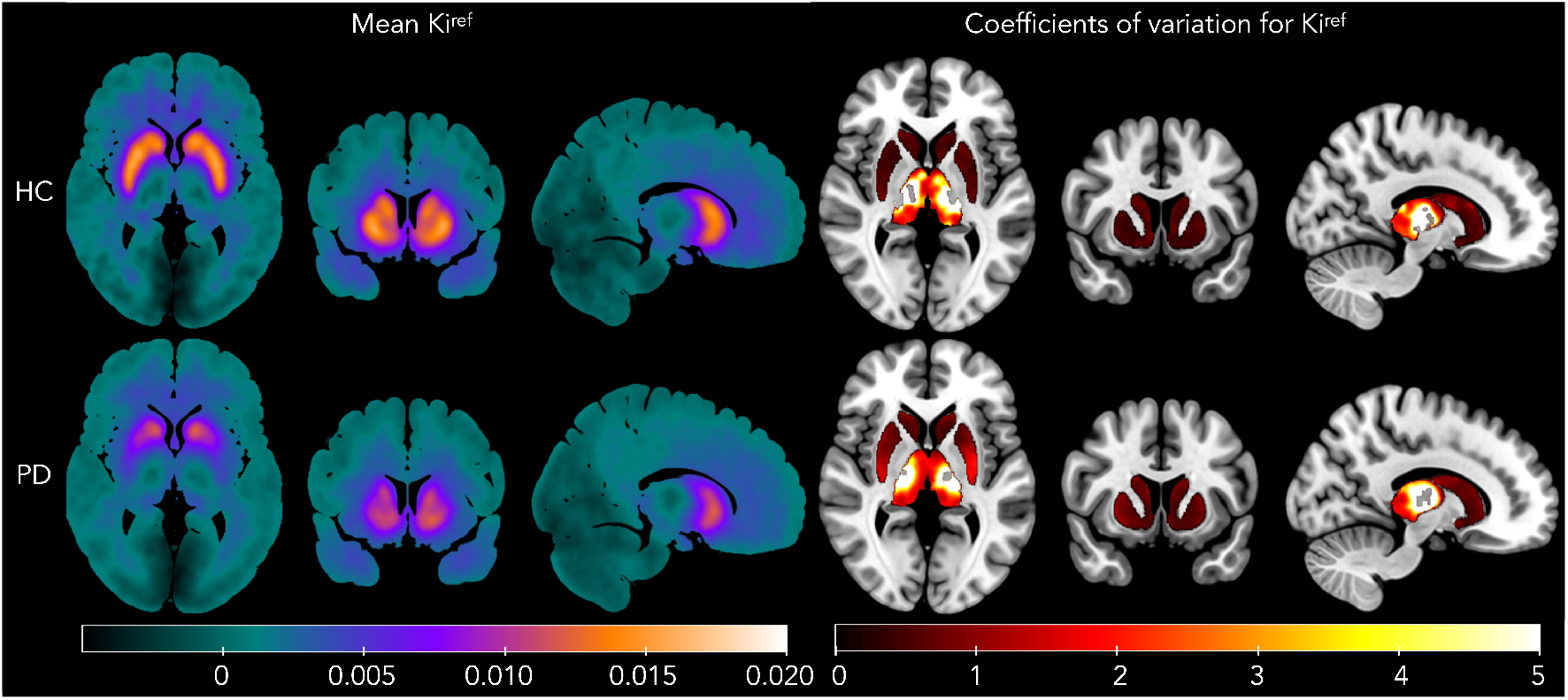
Maps of the Ki^ref^ estimates across healthy controls (HC) and Parkinson’s disease patients (PD), using images normalized on MNI space. Left: Mean Ki^ref^. Right: Coefficients of variation (standard deviation / mean) for Ki^ref^ within the regions of interest (Harvard-Oxford atlas masks). The maps are presented on an MNI template.

**Figure 4.**
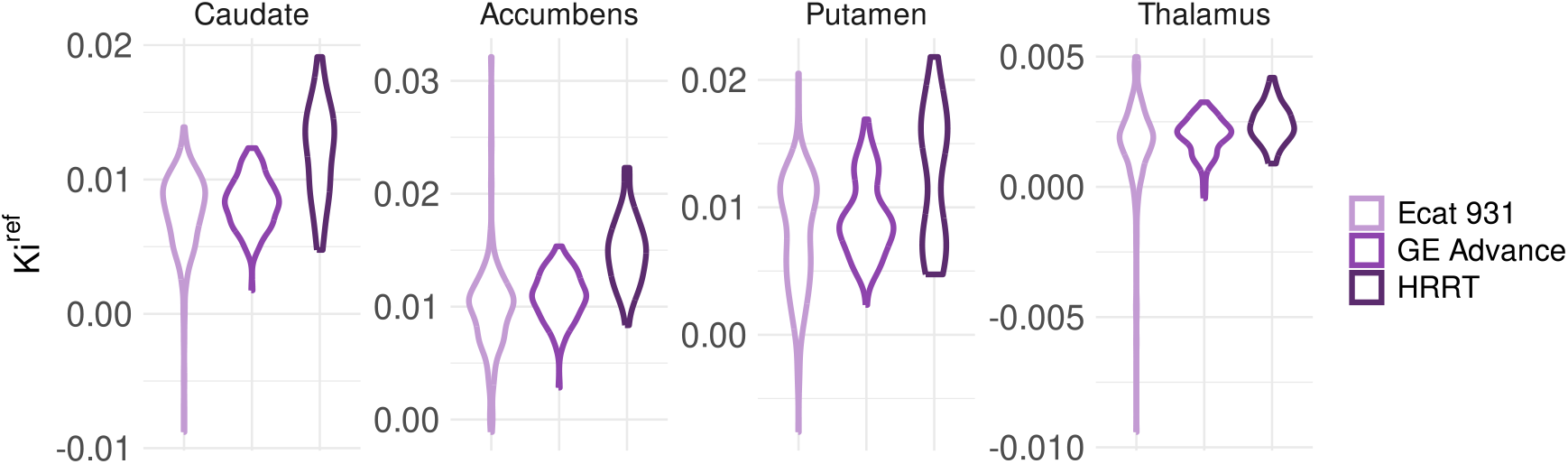
Distributions of scanner-specific Ki^ref^ estimates in the total sample.

### The effects of demographic factors, PD, and scanner

Regression models were used to estimate the effects of age, sex, BMI, PD, and scanner on Ki^ref^. The findings from the three model types – i) effects of age, sex, PD, and scanner in the total sample, ii) the same effects in the BMI subsample, and iii) the same effects and the additional BMI effect in the BMI subsample – were essentially the same.

Ki^ref^ was lower in PD patients compared to healthy controls across striatal ROIs. Ki^ref^ also declined consistently with increasing age, and females had higher Ki^ref^ than males. Finally, Ki^ref^ estimates were systematically higher with newer scanners (GE Advance and particularly HRRT) compared to the reference scanner (Ecat 931).

In the BMI subsample, these effects were amplified (females > males, PD < controls, Ecat 931 < GE Advance < HRRT). Notably, the PD effect in the thalamus differed between analyses: positive in the total sample (PD > controls) but negative in the BMI subsample (PD < controls) with both model types (with and without the BMI effect). In addition, BMI showed a modest positive association with striatal Ki^ref^. This discrepancy likely reflects sample composition. Of the 130 subjects excluded from the BMI subsample, 116 (89%) were scanned with the oldest device (Ecat 931). Thus, the BMI subsample was enriched for higher-quality data (only 30% scanned with Ecat 931), reducing scanner-related noise. Adding BMI as a predictor did not materially alter PD effects.

We therefore considered the BMI subsample model including BMI effect as the most reliable and present its results in **Table 4** and **Figure 5**. Results from the other models are provided in the supplementary material (section **The effects of age, sex, PD, and scanner in the total sample and their validation in the BMI subsample**).

**Table 4.**
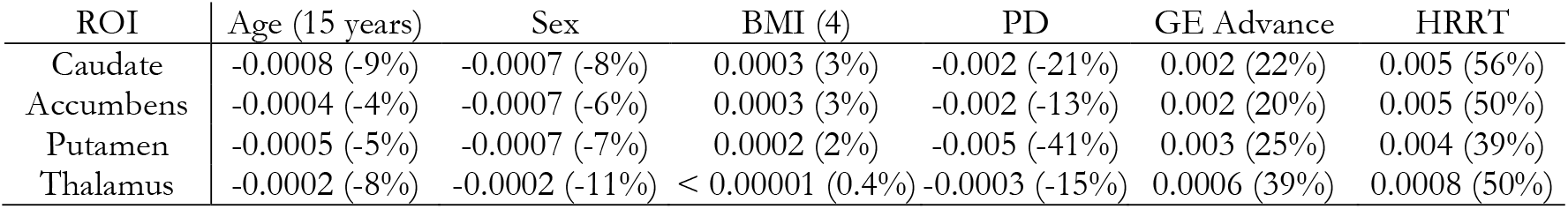
Effect sizes for age (per 15 years), sex (males versus females), BMI (per 4-unit increase), PD (patients versus controls), and scanner type (versus Ecat 931) on regional Ki^ref^, expressed on the original scale. In parenthesis: For the continuous effects of age and BMI, the effect size is given as a percentage of mean regional Ki^ref^ across the sample. For the categorical effects of sex, PD, GE Advance, and HRRT, the effect size is given as a percentage of mean regional Ki^ref^ in the reference category (females, healthy controls, Ecat 931, and Ecat 931 respectively).

**Figure 5.**
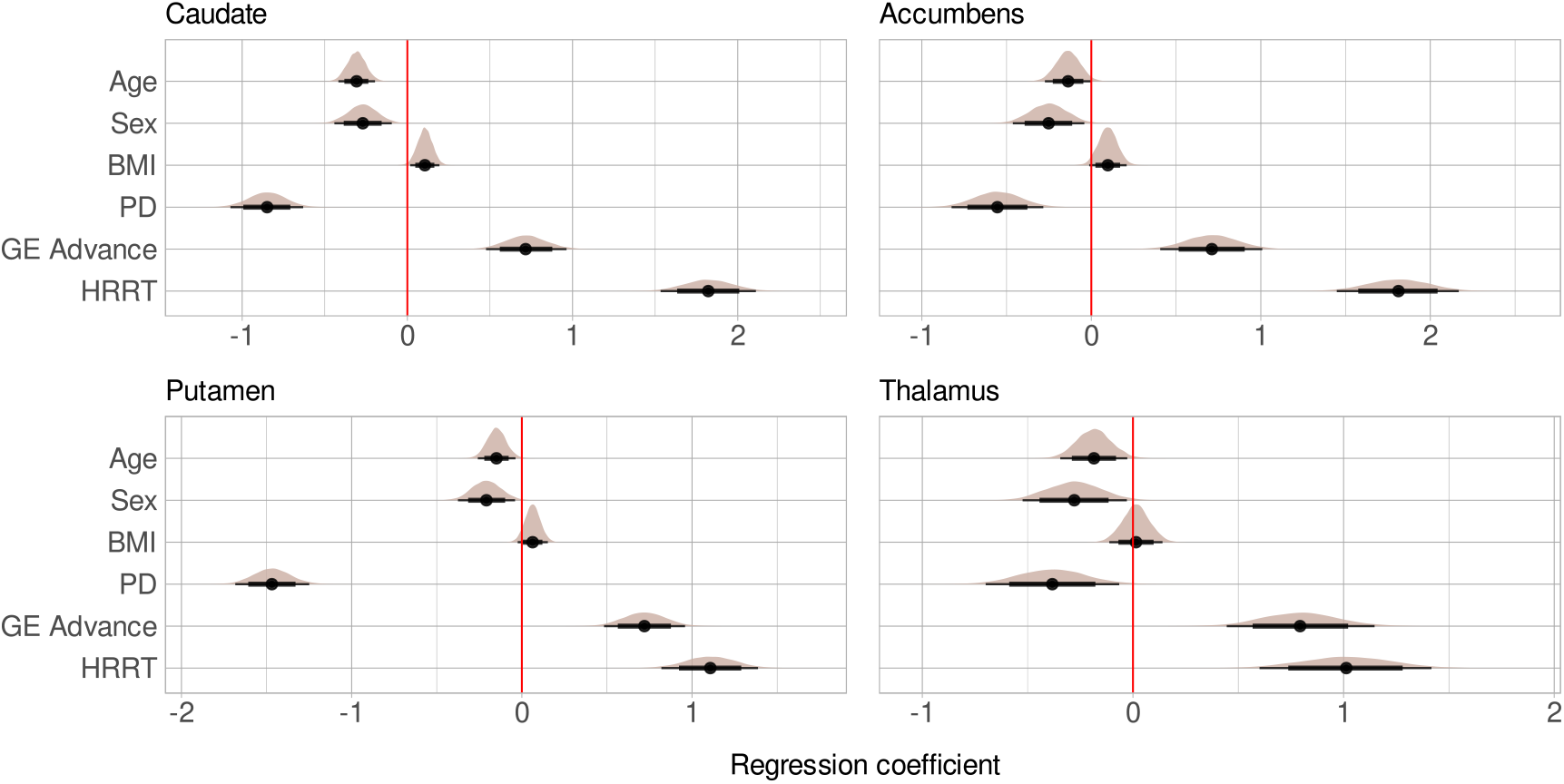
Main effects of age, sex (males versus females), BMI, PD (PD patients versus healthy controls), and scanner (each scanner versus Ecat 931) on Ki^ref^ in the BMI subsample. The figure shows posterior distributions with medians (point) and intervals (95% thin and 80% thick line) on a standardized scale.

### Interaction of age and sex

We assessed the age-effect separately in males and females (see details in supplementary material section **Interaction of age and sex**). These models suggested that the effect of age on Ki^ref^ was negative and roughly of a same magnitude in males and females (comparable to the observed age effects estimated for the males and females together). In accumbens, the negative age-effect was observed in females but not in males.

### Linearity assessment of the age and BMI effects

We estimated the possible nonlinearity of the continuous effects of age and BMI (see details in supplementary material section **Linearity assessment of the age and BMI effects**). Due to uncertainty indicated by wide posterior intervals in the modeling estimates, the possible nonlinearity was difficult to assess, particularly at the ends of the age and BMI ranges. However, there was subtle support for nonlinear relationship between age and Ki^ref^ particularly in caudate and accumbens, suggesting a positive or nonexistent effect until the age of around 50 and a negative effect onwards. We did not see clear support for nonlinear BMI effect in any ROI.

### Interregional correlation of Ki^ref^

The interregional Ki^ref^ Pearson correlations separately in healthy controls and PD patients within each scanner are given with sample sizes in **Figure 6**. Based on Pearson correlation, particularly with the newer scanners HRRT and GE Advance, Ki^ref^ estimates were positively correlated across ROIs in both healthy controls and PD patients.

**Figure 6.**
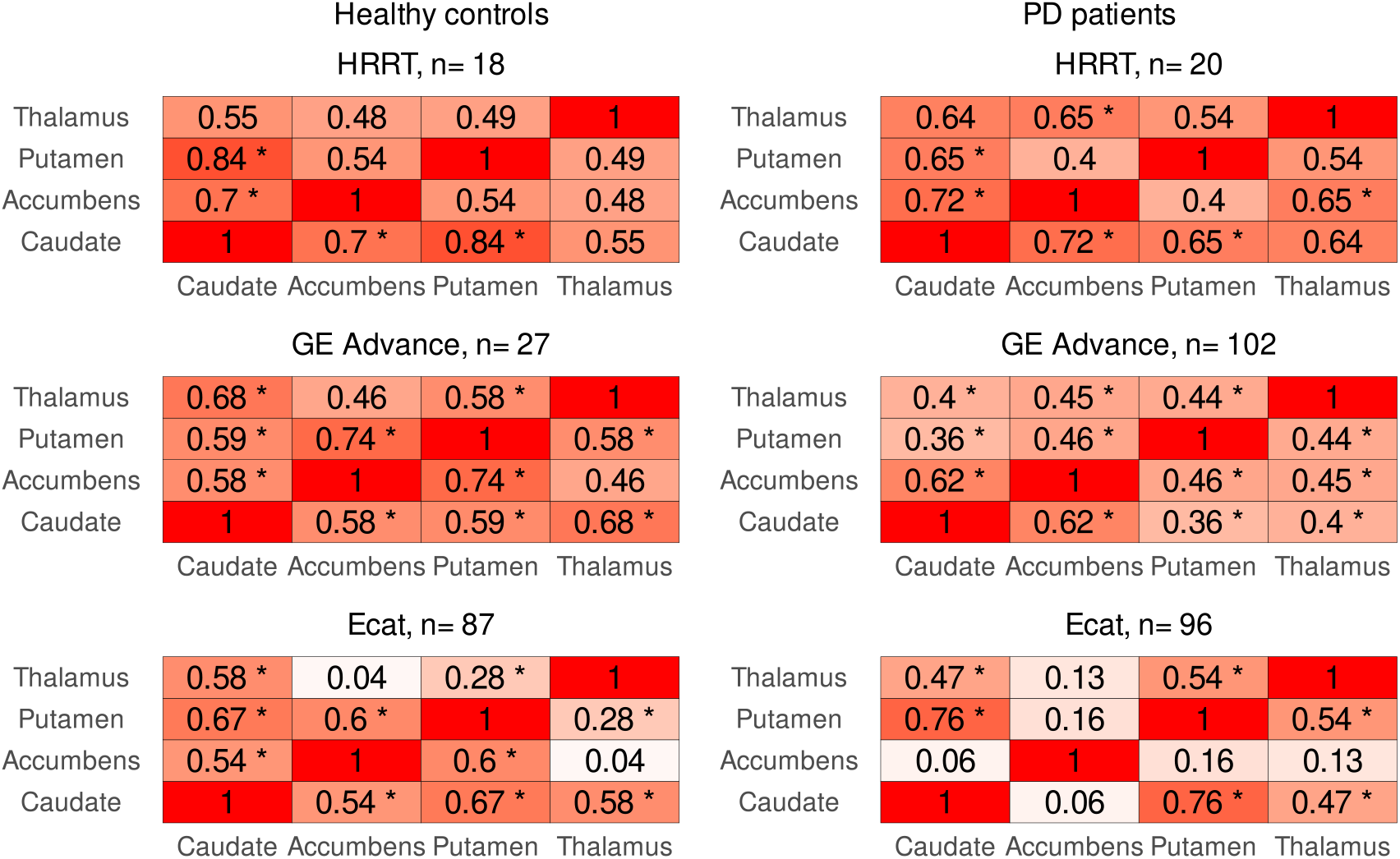
Interregional Ki^ref^ Pearson correlations with sample sizes (n). The table shows point estimates and the p-values of < 0.01 (*). PD= patients with Parkinson’s disease.

### Relationship between striatal dopamine synthesis capacity and the D_2_R availability

The comparison of the [^18^F]fluorodopa Ki^ref^ and [^11^C]raclopride BPND is shown in **Figure 7**. Based on the figure and the Pearson correlation analyses, we did not observe clear support for overall link between regional Ki^ref^ and BPND, although the positive correlation was statistically significant in caudate. The correlation was 0.48 (p= 0.006) in caudate, 0.25 (p= 0.16) in accumbens, 0.013 (p= 0.94) in putamen, and -0.007 (p= 0.97) in thalamus. Given the small sample size, especially for healthy controls, these results should be interpreted with caution.

**Figure 7.**
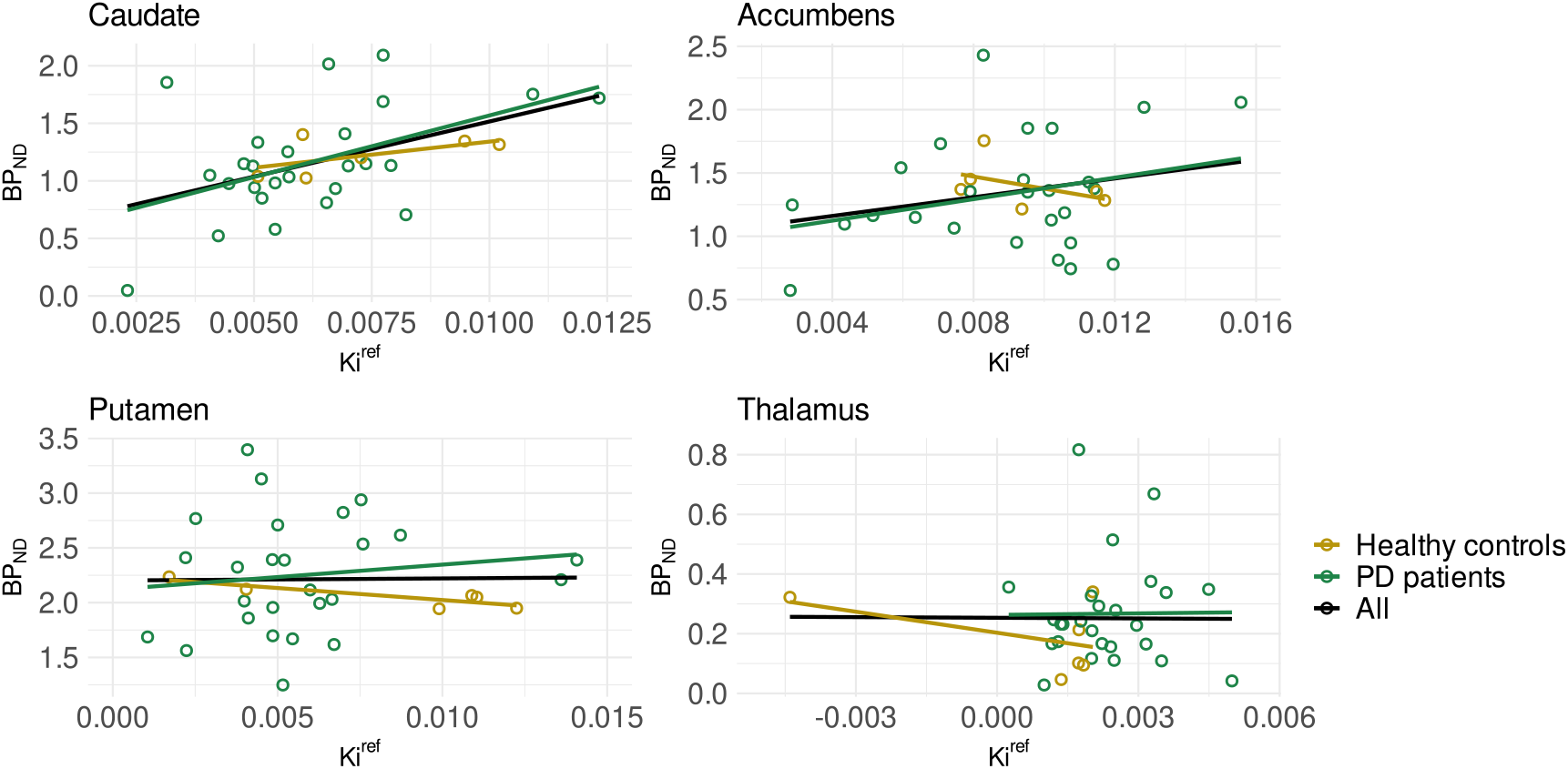
Association between the [^18^F]fluorodopa Ki^ref^ and [^11^C]raclopride BPND in 32 subjects (6 healthy controls, 26 PD patients) in each ROI. The colored lines represent linear fits separately and together for the subject groups.

## Discussion

As expected, PD patients had lower dopamine synthesis capacity compared to healthy controls in the striatum and thalamus. Synthesis capacity also declined through age (all ROIs), increased as a function of BMI (striatal ROIs), and the capacity was higher for females than males. The dopamine synthesis capacity was positively correlated across regions both in healthy controls and in PD patients. Only in caudate, there was support for a positive correlation between dopamine synthesis capacity and D_2_R availability when PD patients and healthy subjects were assessed together. Based on our data involving three different PET cameras, the scanner induces variation in the data, which should be adjusted for when analyzing Ki^ref^ estimates from different devices. Atlas- and MRI-based normalization methods however produced comparable Ki^ref^ estimates for most subjects.

### Dopamine synthesis capacity, Parkinson’s disease and demographic factors

Overall, the dopamine synthesis capacity was substantially lower in PD patients versus healthy controls, and the effect was greater than the demographic effects of age, sex, and BMI. Effect of PD on dopamine synthesis was particularly prominent in the striatal ROIs. The PD related decline in the striatal dopamine synthesis is undebated (Kaasinen & Vahlberg, 2017), yet our findings highlight that the magnitude of the PD effect measurably exceeds the demographic effects. Dopamine synthesis capacity decreased with age both in males and females, yet our data suggested that the decline may appear not until later adulthood (approximately at the age of 50), possibly contributing to cognitive ageing (Karrer et al., 2017). The nonlinearity of the age effect (positive or nonexistent until negative in later adulthood) may have been overlooked in prior studies concluding correlation-based sex-dependency (Laakso et al., 2002), absence, or inconsistency of the age effect (Karrer et al., 2017). Overall, females had higher dopamine synthesis capacity than males in the striatum and thalamus, in line with an earlier analysis with possibly dependent, yet smaller sample from the site (Laakso et al., 2002). Consistently, in a study with [^11^C]raclopride PET, females showed lower D_2_R affinity than males, concluded to support higher endogenous striatal dopamine concentration in females versus males (Pohjalainen et al., 1998).

A recent qualitative review (Janssen & Horstmann, 2022) supported negative association of BMI and dopamine synthesis in striatal subregions, based on three partial correlation studies with the sample sizes of 60 (Lee et al., 2018), 16 (Wallace et al., 2014), and 15 (Wilcox et al., 2010). We utilized these findings in our additional analysis by applying an alternative prior placing higher probability for negative than positive association between BMI and dopamine synthesis capacity. However, even with the alternative prior, our sample of two hundred subjects showed subtle support for a positive (although not sure if linear throughout the BMI range of 18-44) link between dopamine synthesis capacity and BMI in the striatal regions, while no link in thalamus. Taken these conflicting findings together, the relationship between BMI and dopamine synthesis capacity does not appear strong.

### Interregional correlations in dopamine synthesis capacity

To our knowledge, this is the first study to establish significant interregional correlations of dopamine synthesis capacity within striatal and thalamic regions in these groups. High correlation coefficients indicate that individuals with relatively low Ki^ref^ in one region also showed low Ki^ref^ in the remaining regions. These findings, thus, suggest that, in this mixed cohort, also PD is associated with a relatively uniform reduction in dopamine synthesis capacity across striatal subregions. However, because detailed information on disease stage was not systematically available, it is possible that very early PD shows considerably more regionally selective vulnerability.

### Coupling of presynaptic dopamine synthesis and postsynaptic D_2_R

In our dual-tracer subsample (n=32), which consisted mainly of PD patients (81%) and was largely representative of later-stage disease, we found a positive correlation between dopamine synthesis capacity and D_2_R availability in caudate. This pattern suggests that, at advanced stages, presynaptic and postsynaptic dopaminergic markers decline in parallel. Previous studies indicate that the relationship between these markers is dynamic and stage-dependent. In early PD, reduced dopamine synthesis capacity associates with compensatory upregulation of D_2_Rs (Kaasinen et al., 2021). Such upregulation is transients, typically lasting for the first four years after motor symptoms onset (Kaasinen et al., 2021). With progression, both synthesis capacity and receptor availability decline (Antonini et al., 1995), while pharmacological treatments may further explain the correlations through increased competition with endogenous dopamine (Thobois et al., 2004). Our findings therefore align with prior evidence that the compensatory D_2_R upregulation observed in early PD gives way to a phase of coupled presynaptic and postsynaptic degeneration in later disease stages. However, the small sample size and imbalance between patients and controls precule definitive conclusions. Larger stage-stratified studies are required to clarify the trajectory of this relationship across the disease course.

### Scanner affects the dopamine synthesis capacity measures using [^18^F]fluorodopa PET

Scanner contributed to the regional measures of the dopamine synthesis capacity using [^18^F]fluorodopa PET. The device order from highest to lowest Ki^ref^ estimates was HRRT > GE Advance > Ecat 931. It is recommended to adjust for the scanner effect when analyzing data acquired with different scanners. Here, we adjusted for the scanner by estimating the effects of interest independently of the scanner and the interregional correlation within each scanner. There are also other methods, such as data harmonization by scanner (Fortin et al., 2017; Lövdal et al., 2025). However, if the subject-related data is highly dependent on the device (e.g. all patients scanned with one device and healthy controls with another), careful consideration on how to adjust for the scanner is required to avoid confounds (McElreath, 2020).

### Atlas-based spatial normalization can be used with caution to avoid great data loss due to missing MRIs

When individual-level structural MRI is not available, atlas-based normalization provides Ki^ref^ estimates largely comparable with those from MRI-based method for most subjects. The noise could possibly be reduced by adjusting for the scanner, which was not possible here due to limited sample size. Although MRI-based normalization should remain the preferred approach, the atlas-based normalization may be worth applying to avoid great data loss due to missing MRIs, particularly when the sample size is sufficient to cover the possible noise in the data. Nevertheless, whenever both MRI- and atlas-based estimates are available, their agreement should be evaluated to ensure the robustness of findings in future datasets.

## Limitations

The dual-tracer subsample used to examine the relationship between dopamine synthesis capacity and D_2_R availability was small (n= 32) and predominantly composed of PD patients, limiting generalizability. Clinical information, such as disease duration, medication status, and symptom severity was not systematically available. The retrospective dataset spanned three decades and multiple PET scanners, introducing technical heterogeneity despite statistical adjustment. Adjusting for the scanner type could potentially have decreased the noise in the comparison of the spatial normalization methods, yet our sample was limited for such adjustment. An alternative prior assigned in the regression modeling for the effect of sex was based on a sample (Laakso et al., 2002) that possibly, yet only partially, overlapped with our data. However, we made our primary conclusions from models that were not using the alternative prior. Finally, most Ki^ref^ estimates were based on atlas-based normalization rather than subject-specific MRI, which may have added noise to the measurements.

## Conclusions

PD significantly reduces striatal dopamine synthesis capacity. Demographic factors also influence synthesis capacity independently of PD, but these effects are smaller than those accounted by PD. Synthesis capacity declined with aging and particularly after the age of ∼50, was higher in females than males, and showed a possible association with BMI in striatal regions. Although thalamic findings were uncertain, they were mainly consistent with the overall pattern in the striatum.

The dopamine synthesis capacity was strongly correlated across striatal and thalamic regions in both healthy controls and PD patients, suggesting a relatively uniform interregional effect. In a limited dual-tracer subsample that likely overrepresented late-stage PD, we observed a positive correlation between dopamine synthesis capacity and D_2_R availability in caudate, yet no clear associations in the other regions.

Methodologically, scanner had a substantial influence on Ki^ref^ estimates, emphasizing the need to adjust for scanner effects in multi-scanners datasets. When MRI is unavailable, atlas-based normalization may be used to prevent data loss, though MRI-based normalization should remain the preferred approach.

## Data availability

The mean brain maps of group specific (healthy controls and PD patients) dopamine synthesis capacities are available in NeuroVault (https://neurovault.org/collections/OKUNGLWA/).

## Acknowledgements

This study was supported by the Finnish Cultural Foundation (grants to T.M.), the Finnish Governmental Research Funding for Turku University Hospital and for the Western Finland collaborative area, the Päivikki and Sakari Sohlberg Foundation, and the Finnish Brain Foundation. We acknowledge Senior Research Fellow and modeling specialist Vesa Oikonen for consulting on the Patlak modeling, as well as Professor Sven De Maeyer from the University of Antwerp, Belgium, and data engineer and scientist Tomi Karjalainen for their expertise in the Bayesian regression modeling and visualization. We thank Tuomas Knuuti, Lilli Laine, Veera Soininen, and Jenny Lindman for conducting quality control for the preprocessed imaging data.

