## Supplementary material for "Striatal dopamine synthesis capacity in Parkinson’s disease: Effects of age, sex, and body mass index in a large [^18^F]fluorodopa PET cohort"

**Publications whose data are used in the current study** (might not be comprehensive)

Although we did not have systematic information of the specific publications where each [$^{18}$F]fluorodopa PET image in our data was originally reported, we here list the Turku PET Centre publications that we are aware of from the imaging years 1988-2016 (+4 years) potentially reporting the same [$^{18}$F]fluorodopa data of healthy controls and/ or Parkinson's disease (PD) patients than here.

Rinne, J., Laihinen, A., Någren, K., Bergman, J., Solin, O., Haaparanta, M., Ruotsalainen, U., & Rinne, U. (1990). PET demonstrates different behaviour of striatal dopamine D-1 and D-2 receptors in early Parkinson's disease. *Journal of neuroscience research*, *27*(4), 494-499. (Rinne et al., 1990).

Laihinen, A., Rinne, J., Rinne, U., Haaparanta, M., Ruotsalainen, U., Bergman, J., & Solin, O. (1992). [18F]-6-fluorodopa PET scanning in Parkinson's disease after selective COMT inhibition with nitecapone (OR-462). *Neurology*, *42*(1), 199-199. (Laihinen et al., 1992).

Hietala, J., Syvälahti, E., Vuorio, K., Räkköläinen, V., Bergman, J., Haaparanta, M., Solin, O., Kuoppamäki, M., Kirvelä, O., & Ruotsalainen, U. (1995). Presynaptic dopamine function in striatum of neuroleptic-naive schizophrenic patients. *Lancet (London, England)*, *346*(8983), 1130-1131. (Hietala et al., 1995).

Ruottinen, H., Rinne, J., Ruotsalainen, U., Bergman, J., Oikonen, V., Haaparanta, M., Solin, O., Laihinen, A., & Rinne, U. (1995). Striatal [18 F] fluorodopa utilization after COMT inhibition with entacapone studied with PET in advanced Parkinson's disease. *Journal of neural transmission-Parkinson's disease and dementia section*, *10*, 91-106. (Ruottinen et al., 1995).

Ruottinen, H. M., Rinne, J. O., Oikonen, V. J., Bergman, J. R., Haaparanta, M. T., Solin, O. H., Ruotsalainen, U., & Rinne, U. K. (1997). Striatal 6-[18F] fluorodopa accumulation after combined inhibition of peripheral catechol-O-methyltransferase and monoamine oxidase type B: Differing response in relation to presynaptic dopaminergic dysfunction. *Synapse*, *27*(4), 336-346. (Ruottinen et al., 1997).

Tiihonen, J., Vilkman, H., Räsänen, P., Ryynänen, O., Hakko, H., Bergman, J., Hämäläinen, T, Laakso, A., Haaparanta-Solin, M., & Solin, O. (1998). Striatal presynaptic dopamine function in type 1 alcoholics measured with positron emission tomography. *Molecular Psychiatry*, *3*(2), 156-161. (Tiihonen et al., 1998).

Hietala, J., Syvälahti, E., Vilkman, H., Vuorio, K., Räkköläinen, V., Bergman, J., Haaparanta, M., Solin, O., Kuoppamäki, M., Eronen, E., Ruotsalainen, U., & Salokangas, R. K. (1999). Depressive symptoms and presynaptic dopamine function in neuroleptic-naive schizophrenia. *Schizophrenia research*, *35*(1), 41-50. (Hietala et al., 1999).

Laihinen, A., Ruottinen, H., Rinne, J., Haaparanta, M., Bergman, J., Solin, O., Koskenvuo, M., Marttila, R., & Rinne, U. (2000). Risk for Parkinson's disease: twin studies for the detection of asymptomatic subjects using [18 F] 6-fluorodopa PET. *Journal of Neurology*, *247*, 110-113. (Laihinen et al., 2000).

Rinne, J. O., Portin, R., Ruottinen, H., Nurmi, E., Bergman, J., Haaparanta, M., & Solin, O. (2000). Cognitive impairment and the brain dopaminergic system in Parkinson disease:[18F] fluorodopa positron emission tomographic study. *Archives of Neurology*, *57*(4), 470-475. (Rinne et al., 2000).

Salokangas, R. K., Vilkman, H., Ilonen, T., Taiminen, T., Bergman, J., Haaparanta, M., Solin, O., Alanen, A., Syvälahti, E., & Hietala, J. (2000). High levels of dopamine activity in the basal ganglia of cigarette smokers. *American Journal of Psychiatry*, *157*(4), 632-634. (Salokangas et al., 2000).

Brück, A., Portin, R., Lindell, A., Laihinen, A., Bergman, J., Haaparanta, M., Solin, O., & Rinne, J. (2001). PET shows that impaired frontal lobe functioning in Parkinson's disease is related to dopaminergic hypofunction in the caudate nucleus. *Neuroscience letters*, *311*, 81-84. (Brück et al., 2001).

Kaasinen, V., Nurmi, E., Bergman, J., Eskola, O., Solin, O., Sonninen, P., & Rinne, J. O. (2001). Personality traits and brain dopaminergic function in Parkinson's disease. *Proceedings of the National Academy of Sciences*, *98*(23), 13272-13277. (Kaasinen, Nurmi, Bergman, et al., 2001).

Kaasinen, V., Nurmi, E., Brück, A., Eskola, O., Bergman, J., Solin, O., & Rinne, J. (2001). Increased frontal [18F] fluorodopa uptake in early Parkinson's disease: sex differences in the prefrontal cortex. *Brain*, *124*(6), 1125-1130. (Kaasinen, Nurmi, Brück, et al., 2001).

Nurmi, E., Ruottinen, H. M., Bergman, J., Haaparanta, M., Solin, O., Sonninen, P., & Rinne, J. O. (2001). Rate of progression in Parkinson's disease: a 6-[18F] fluoro-L-dopa PET study. *Movement disorders: official journal of the Movement Disorder Society*, *16*(4), 608-615. (Nurmi et al., 2001).

Rinne, J. O., Nurmi, E., Ruottinen, H. M., Bergman, J., Eskola, O., & Solin, O. (2001). [18F] FDOPA and [18F] CFT are both sensitive PET markers to detect presynaptic dopaminergic hypofunction in early Parkinson's disease. *Synapse*, *40*(3), 193-200. (Rinne et al., 2001).

Ruottinen, H. M., Niinivirta, M., Bergman, J., Oikonen, V., Solin, O., Eskola, O., Eronen, E., Sonninen, P., & Rinne, U. K. (2001). Detection of response to COMT inhibition in FDOPA PET in advanced Parkinson's disease requires prolonged imaging. *Synapse*, *40*(1), 19-26. (Ruottinen et al., 2001).

Kaasinen, V., Nurmi, E., Bergman, J., Solin, O., Kurki, T., & Rinne, J. O. (2002). Personality traits and striatal 6-[18F] fluoro-L-dopa uptake in healthy elderly subjects. *Neuroscience letters*, *332*(1), 61-64. (Kaasinen et al., 2002).

Laakso, A., Vilkman, H., Örgen Bergman, J., Haaparanta, M., Solin, O., Syvälahti, E., Salokangas, R. K., & Hietala, J. (2002). Sex differences in striatal presynaptic dopamine synthesis capacity in healthy subjects. *Biological psychiatry*, *52*(7), 759-763. (Laakso et al., 2002).

Laakso, A., Wallius, E., Kajander, J., Bergman, J., Eskola, O., Solin, O., Ilonen, T., Salokangas, R. K., Syvälahti, E., & Hietala, J. (2003). Personality traits and striatal dopamine synthesis capacity in healthy subjects. *American Journal of Psychiatry*, *160*(5), 904-910. (Laakso et al., 2003).

Brück, A., Aalto, S., Nurmi, E., Bergman, J., & Rinne, J. O. (2005). Cortical 6-[18F] fluoro-L-dopa uptake and frontal cognitive functions in early Parkinson's disease. *Neurobiology of aging*, *26*(6), 891-898. (Brück et al., 2005).

Laakso, A., Pohjalainen, T., Bergman, J., Kajander, J., Haaparanta, M., Solin, O., Syvälahti, E., & Hietala, J. (2005). The A1 allele of the human D2 dopamine receptor gene is associated with increased activity of striatal L-amino acid decarboxylase in healthy subjects. *Pharmacogenetics and genomics*, *15*(6), 387-391. (Laakso et al., 2005).

Brück, A., Aalto, S., Nurmi, E., Vahlberg, T., Bergman, J., & Rinne, J. O. (2006). Striatal subregional 6-[18F] fluoro-L-dopa uptake in early Parkinson's disease: A two-year follow-up study. *Movement disorders: official journal of the Movement Disorder Society*, *21*(7), 958-963. (Brück et al., 2006).

Huttunen, J., Heinimaa, M., Svirskis, T., Nyman, M., Kajander, J., Forsback, S., Solin, O., Ilonen, T., Korkeila, J., & Ristkari, T. (2008). Striatal dopamine synthesis in first-degree relatives of patients with schizophrenia. *Biological psychiatry*, *63*(1), 114-117. (Huttunen et al., 2008).

Brück, A., Aalto, S., Rauhala, E., Bergman, J., Marttila, R., & Rinne, J. O. (2009). A follow-up study on 6-[18F] fluoro-L-dopa uptake in early Parkinson's disease shows nonlinear progression in the putamen. *Movement Disorders*, *24*(7), 1009-1015. (Brück et al., 2009).

Jokinen, P., Brück, A., Aalto, S., Forsback, S., Parkkola, R., & Rinne, J. O. (2009). Impaired cognitive performance in Parkinson's disease is related to caudate dopaminergic hypofunction and hippocampal atrophy. *Parkinsonism & related disorders*, *15*(2), 88-93. (Jokinen, Brück, et al., 2009).

Jokinen, P., Helenius, H., Rauhala, E., Brück, A., Eskola, O., & Rinne, J. O. (2009). Simple ratio analysis of 18F-fluorodopa uptake in striatal subregions separates patients with early Parkinson disease from healthy controls. *Journal of Nuclear Medicine*, *50*(6), 893-899. (Jokinen, Helenius, et al., 2009).

Joutsa, J., Martikainen, K., Niemelä, S., Johansson, J., Forsback, S., Rinne, J. O., & Kaasinen, V. (2012). Increased medial orbitofrontal [18F] fluorodopa uptake in Parkinsonian impulse control disorders. *Movement Disorders*, *27*(6), 778-782. (Joutsa et al., 2012).

Kaasinen, V., Jokinen, P., Joutsa, J., Eskola, O., & Rinne, J. O. (2012). Seasonality of striatal dopamine synthesis capacity in Parkinson's disease. *Neuroscience letters*, *530*(1), 80-84. (Kaasinen et al., 2012).

Jokinen, P., Karrasch, M., Brück, A., Johansson, J., Bergman, J., & Rinne, J. O. (2013). Cognitive slowing in Parkinson's disease is related to frontostriatal dopaminergic dysfunction. *Journal of the neurological sciences*, *329*(1-2), 23-28. (Jokinen et al., 2013).

Joutsa, J., Rinne, J. O., Eskola, O., & Kaasinen, V. (2013). Reduced striatal dopamine synthesis capacity is associated with symptoms of depression in patients with de novo unmedicated Parkinson's disease. *Journal of Parkinson's Disease*, *3*(3), 325-329. (Joutsa et al., 2013).

Järvelä, J. T., Rinne, J. O., Eskola, O., & Kaasinen, V. (2014). Mortality in Parkinson's disease is not associated with the severity of early dopaminergic defect. *Parkinsonism & related disorders*, *20*(8), 894-897. (Järvelä et al., 2014).

Joutsa, J., Voon, V., Johansson, J., Niemelä, S., Bergman, J., & Kaasinen, V. (2015). Dopaminergic function and intertemporal choice. *Translational psychiatry*, *5*(1), e491-e491. (Joutsa et al., 2015).

Majuri, J., Joutsa, J., Johansson, J., Voon, V., Alakurtti, K., Parkkola, R., Lahti, T., Alho, H., Hirvonen, J., & Arponen, E. (2017). Dopamine and opioid neurotransmission in behavioral addictions: a comparative PET study in pathological gambling and binge eating. *Neuropsychopharmacology*, *42*(5), 1169-1177. (Majuri et al., 2017).

Majuri, J., Joutsa, J., Arponen, E., Forsback, S., & Kaasinen, V. (2018). Dopamine synthesis capacity correlates with μ-opioid receptor availability in the human basal ganglia: A triple-tracer PET study. *NeuroImage*, *183*, 1-6. (Majuri et al., 2018).

### Comparison of atlas-based and MRI-based $Ki^{ref}$ estimates

For validation purposes, we compared the $Ki^{ref}$ estimates from two spatial normalization methods using either subject-specific magnetic resonance image (MRI) or common PET atlas in the subjects (n= 45, 13% of the total sample) with the $Ki^{ref}$ estimates from both methods available. We show descriptive statistics of the subsample in **Table S1**.

| Group | Age (years) | | | Sex | | BMI (n= 33) | | | Scanner | | |
|---|---|---|---|---|---|---|---|---|---|---|---|
| | Mean | SD | Range | Male | Female | Mean | SD | Range | Ecat 931 | GE Advance | HRRT |
| HC (n= 22) | 44.4 | 14.0 | 19.5-65.3 | 12 | 10 | 26.0 | 3.9 | 21.8 | 0 | 4 | 18 |
| PD (n= 23) | 63.6 | 9.8 | 33.0-76.4 | 14 | 9 | 26.2 | 6.3 | 38.8 | 0 | 3 | 20 |

**Table S1**. Demographic and imaging characteristics of the subsample of subjects used for comparing the two spatial normalization methods (n= 45). HC= healthy controls, PD= Parkinson's disease patients, SD= standard deviation. Body mass index (BMI) is given for 33 subjects (20 healthy controls and 13 PD patients) with the BMI information available. Scanner column shows the number of subjects imaged with each scanner.

We used correlation analysis to compare the $Ki^{ref}$ estimates from the two methods. We tested the normality of the $Ki^{ref}$ estimates from each of the methods within each ROI using Shapiro-Wilk testing. About a half of the tests (5/8) did not conflict with the null-hypothesis of normally distributed data (significance level 0.01), and we decided to estimate the correlations using both Pearson and rank-order based Spearman correlation. We used one-tailed testing because we expected the possible correlation between the two $Ki^{ref}$ estimates to be positive. Otherwise, we used the default specification of cor.test function in R package stats (R Core Team, 2025). We assessed the normality (Shapiro-Wilk) and the (Pearson and Spearman) correlations using the R package stats (R Core Team, 2025).

The preprocessing methods produced the most comparable $Ki^{ref}$ estimates in caudate and putamen. The one-tailed (greater) Pearson correlation between the $Ki^{ref}$ estimates of the two alternative methods was 0.57 ($p< 0.001$) in caudate, 0.13 ($p= 0.19$) in accumbens, 0.79 ($p< 0.001$) in putamen, and 0.46 ($p< 0.001$) in thalamus. The one-tailed (greater) Spearman rank-order correlation between the $Ki^{ref}$ estimates of the two alternative methods was 0.69 ($p< 0.001$) in caudate, 0.40 ($p= 0.003$) in accumbens, 0.79 ($p< 0.001$) in putamen, and 0.45 ($p= 0.001$) in thalamus. **Figure S1** illustrates the comparativeness of the $Ki^{ref}$ estimates, separately in healthy controls and PD patients.

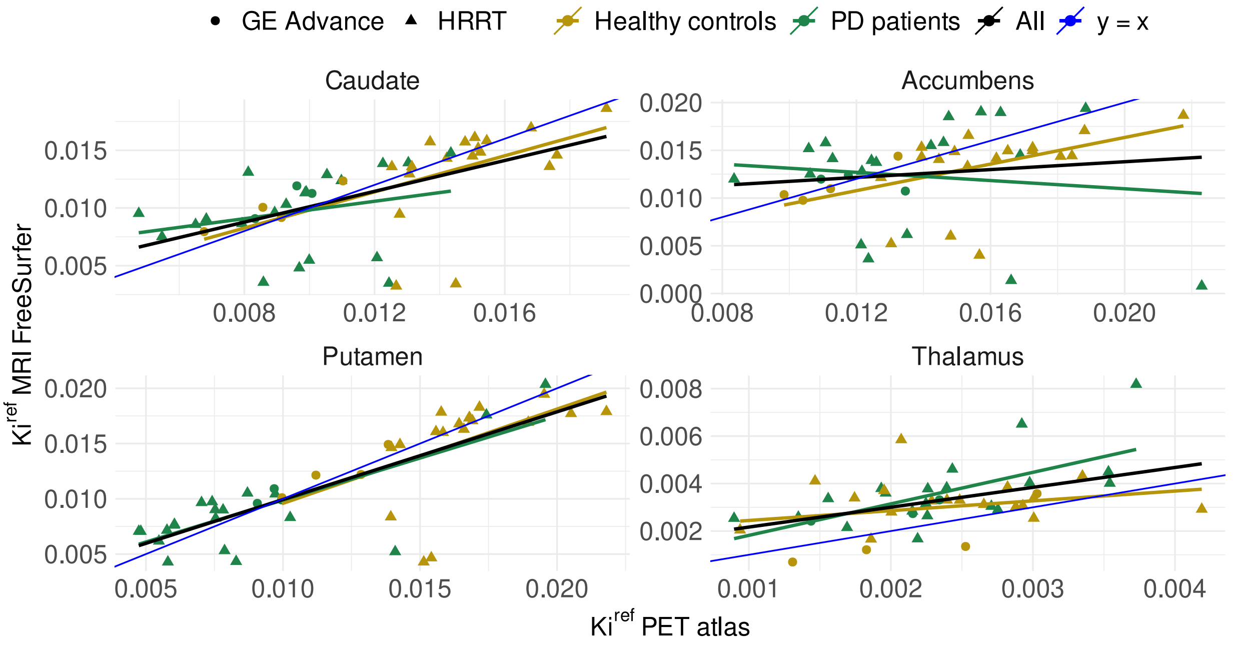


**Figure S1**. $Ki^{ref}$ estimates from the two spatial normalization methods in 45 subjects (22 healthy controls and 23 PD patients) who had the $Ki^{ref}$ from both methods available. The lines represent linear fits separately and together for the subject groups, as well as the y=x line. The individual observations are colored by the group and shaped by the scanner.

Based on the correlation analysis and the **Figure S1**, atlas-based normalization method works rather well for most subjects. Still, there appears noise in the data and thus, the MRI-based normalization is recommended as the primary method. The noise could possibly be reduced by adjusting for the scanner, which was not possible here due to limited sample size. The observations deviating the most from the y = x line are from the scanner HRRT, yet most of the observations (38/45) have been acquired with HRRT. Thus, it is difficult to determine whether the estimates are less comparable with HRRT than GE Advance or any other scanner. However, the atlas-based normalization may be worth using when MRI is not available for the given subject and when the impact of noise can sufficiently be decreased with a large sample size.

### Lateralization of $Ki^{ref}$ estimates

We assessed the lateralization of $Ki^{ref}$ estimates that were spatially normalized using the subject-specific MRI and FreeSurfer. The sample (n= 45) was the same we used for comparing the two normalization methods (**Table S1**). Based on **Figure S2**, it is not systematic which hemisphere has higher $Ki^{ref}$ for either healthy controls or PD patients, and the magnitude of lateralization does not depend on sex. In putamen, there is a clear drop in the overall $Ki^{ref}$ of PD patients versus healthy controls.

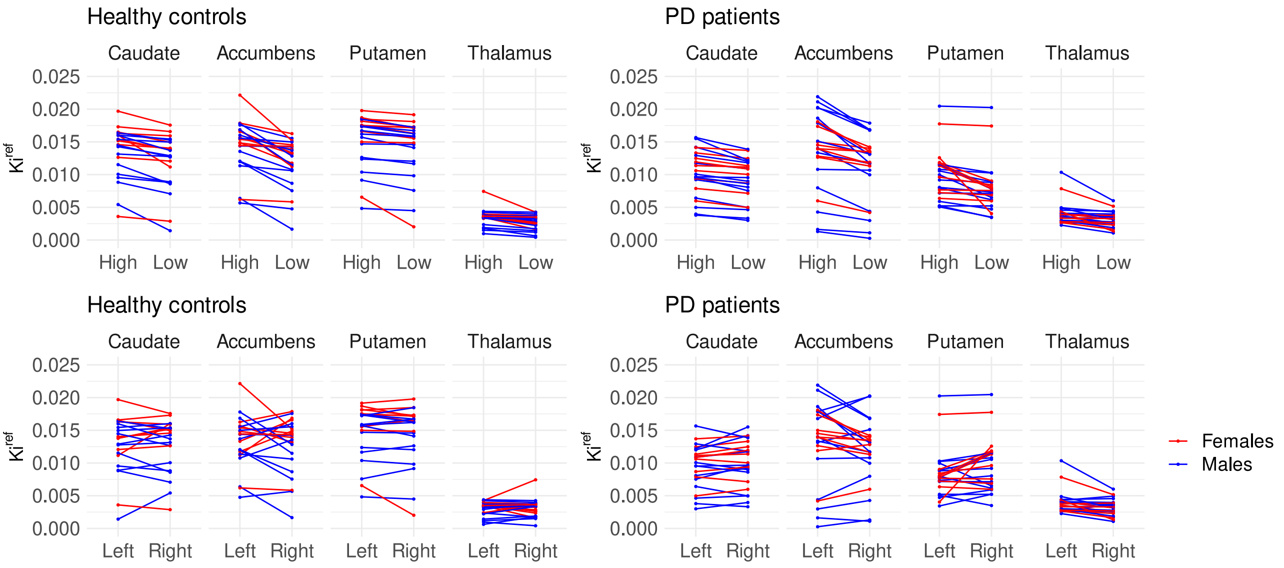


**Figure S2**. The lines join the observations of the hemispheres with higher and lower (top row), as well as left and right (bottom row) $Ki^{ref}$ estimates of each subject.

Asymmetry index was calculated as $[(\text{higher } Ki^{ref} - \text{lower } Ki^{ref})/(\text{higher } Ki^{ref} + \text{lower } Ki^{ref})]*100\%$, where higher refers to the hemisphere with higher $Ki^{ref}$ estimate and lower to the hemisphere with lower $Ki^{ref}$ estimate. Asymmetry indexes are visualized in **Figure S3**. The asymmetry indexes are mainly below twenty for both healthy controls and PD patients, although some outliers exist.

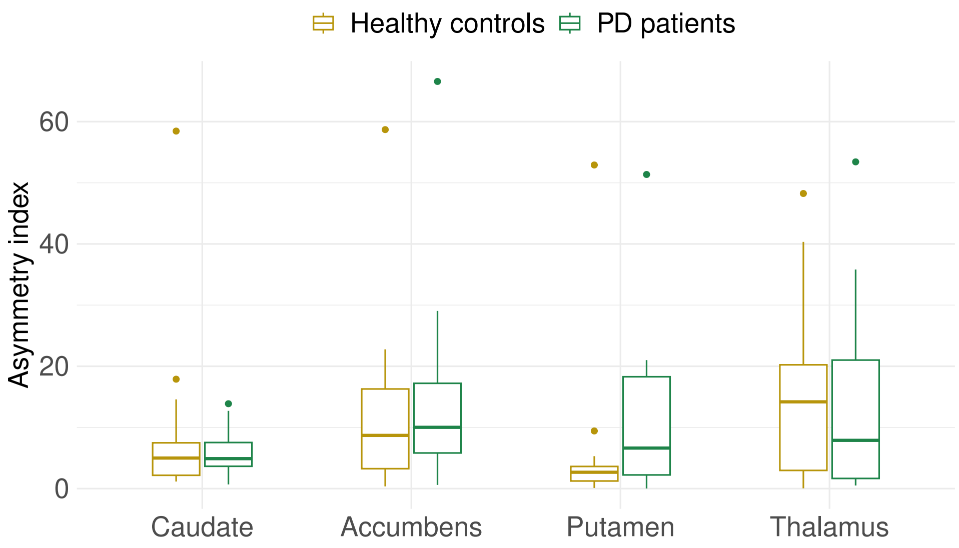


**Figure S3**. Asymmetry indexes as boxplots in healthy controls and PD patients. Each boxplot shows the median (middle line), and 25% (lower hinge) and 75% (upper hinge) quartiles. The lower whisker subtracts 1.5 * IQR from the lower hinge, and the upper whisker adds 1.5 * IQR in the upper hinge. IQR= Inter-quartile range, meaning the distance between the 25% and 75% quartiles. Asymmetry indexes beyond the whiskers are plotted individually.

Using R package stats (R Core Team, 2025), we conducted Shapiro-Wilk normality testing of the asymmetry index in each ROI and separately in healthy controls and PD patients. Most of the tests conflicted with the null hypothesis of normally distributed asymmetry indexes (significance level 0.01), and we had limited number of observations (22 healthy controls, and 23 PD patients). Thus, we carried out nonparametric Wilcoxon rank-sum tests (aka Mann-Whitney test, also using the R package stats) to assess whether the asymmetry indexes differ between healthy controls and PD patients. We used one-sided testing, expecting greater asymmetry in PD patients compared to healthy controls. The testing suggested that in putamen, the asymmetry index might be higher in PD patients than in healthy controls ($p = 0.02$). It is, however, noteworthy, that in putamen there is great variation in the asymmetry indexes of the PD patients compared to the healthy controls (**Figure S3**). The Wilcoxon rank-sum testing did not support the higher asymmetry of PD patients versus healthy controls in the remaining ROIs (the p-value was 0.44 in caudate, 0.23 in accumbens, and 0.73 in thalamus).

Altogether, it appears that some lateralization of the $Ki^{ref}$ estimates exists in both groups but not in all subjects, while the magnitude and direction of the lateralization vary across subjects (**Figure S2**). Furthermore, the asymmetry indexes did not clearly differ between the groups (Wilcoxon and **Figure S3**).

If the potential PD-induced dopamine decline in one hemisphere would be compensated by increased dopamine synthesis in the other, the opposing changes could cancel out and overlook the altered dopamine synthesis capacity in PD compared to healthy controls. However, we do not detect such pattern here. The mean of left and right $Ki^{ref}$ estimates will decrease if the estimate decreases in either of the hemispheres. Thus, we expect that the potential diminished dopamine synthesis in PD is not overlooked when using the mean of left and right $Ki^{ref}$.

**R syntax for the regression models**

Activation of the required R packages:

```
library(rstan)

library(brms)
```

Model specifications:

```
rstan_options(auto_write = TRUE)
ITER <- 4000
WARMUP <- 1000
NUM_CORES <- 4
CHAINS <- 4

# Primary priors:

custom_prior <- c(set_prior('normal(0,1)', class = 'b'))


# Alternative priors:

custom_prior_alt <- c(set_prior('normal(-1,0.5)', class = 'b', coef = c('sexm', 'bmi_z', 'groupparkinson')),
                      set_prior('normal(0,1)', class = 'b', coef = c('age_z', 'scannerGEAdvance',
                      'scannerHRRT')))
```

In the regression modeling, the dependent variable was the standardized regional $Ki^{ref}$ (e.g. cau_kiref_z for caudate). The predictors were standardized age (age_z), sex (male, female), group (healthy control, PD patient), standardized BMI (bmi_z), and scanner (Ecat 931, GE Advance, HRRT). In the syntax, intercept is specified as '1'. In the example syntax given, the region is caudate nucleus (cau), but the same model was estimated for each ROI.

First, we show the syntax for regression modeling of the linear effects of standardized age, sex, standardized BMI, PD, and scanner.

```
fit_cau_bmi <- brm(
  formula= cau_kiref_z ~ 1 + age_z + sex + group + bmi_z + scanner,
  data = BMI subsample,
  prior = custom_prior (and separately using custom_prior_alt),
  cores = NUM_CORES,
  chains = CHAINS,
  iter = ITER,
  warmup = WARMUP,
  sample_prior = TRUE
)
```

Second, we show the syntax for regression modeling of the linear effects of standardized age, sex, PD, and scanner.

```
fit_cau <- brm(
  formula= cau_kiref_z ~ 1 + age_z + sex + group + scanner,
  data = total sample (and separately using BMI subsample),
  prior = custom_prior,
  cores = NUM_CORES,
  chains = CHAINS,
  iter = ITER,
  warmup = WARMUP,
  sample_prior = TRUE
)
```

Third, we show the syntax for regression modeling of the effects of age separately for males and females (interaction of standardized age and sex), standardized BMI, PD, and scanner.

```
fit_cau_bmi_int <- brm(
  formula= cau_kiref_z ~ 1 + age_z*sex + group + bmi_z + scanner,
  data = BMI subsample,
  prior = custom_prior,
  cores = NUM_CORES,
  chains = CHAINS,
  iter = ITER,
  warmup = WARMUP,
  sample_prior = TRUE
)
```

Finally, we show the syntax for regression modeling of the effects of standardized age, sex, standardized BMI, PD, and scanner, while nonlinearity of the continuous effects of standardized age and BMI was allowed. For this model, we specified adapt delta and maximum treedepth.

```
AD <- 0.999
MAX_TREEDEPTH <- 20

fit_cau_bmi_nl <- brm(
  formula= cau_kiref_z ~ 1 + s(age_z) + sex + group + s(bmi_z) + scanner,
  data = BMI subsample,
  prior = custom_prior,
  cores = NUM_CORES,
  chains = CHAINS,
  iter = ITER,
  warmup = WARMUP,
  control = list(adapt_delta = AD, max_treedepth = MAX_TREEDEPTH),
  sample_prior = TRUE
)
```

**The effects of age, sex, PD, and scanner in the total sample and their validation in the BMI subsample**

The results of the initial regression modeling of the effects of age, sex, PD, and scanner on $Ki^{ref}$ in the total sample are given in **Figure S4**.

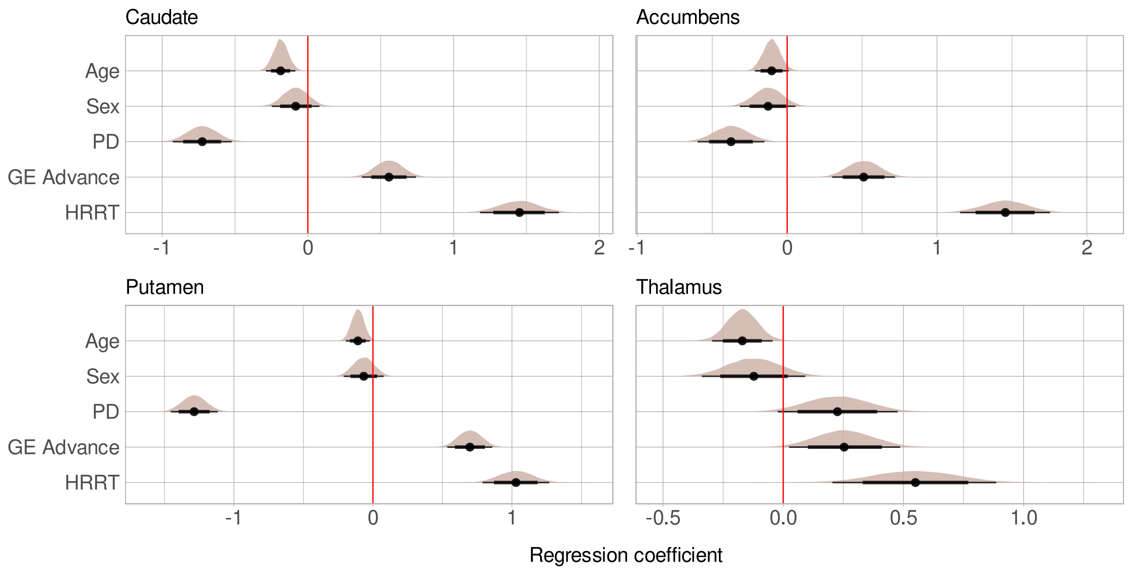


**Figure S4**. Main effects of age, sex (males versus females), PD (PD patients versus healthy controls), and scanner (each scanner versus Ecat 931) on $Ki^{ref}$ in the total sample. The figure shows posterior distributions with medians (point) and intervals (95% thin and 80% thick line) on a standardized scale.

The striatal effects of age, sex, PD, and scanner were validated in the BMI subsample (n= 220). The findings are illustrated in **Figure S5**.

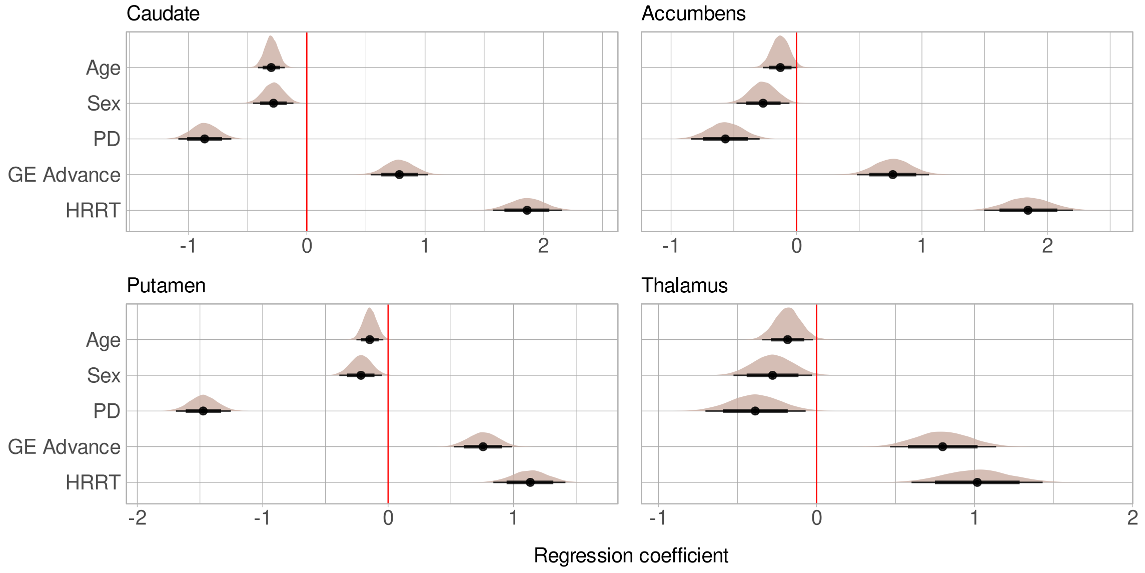


**Figure S5**. Main effects of age, sex (males versus females), PD (PD patients versus healthy controls), and scanner (each scanner versus Ecat 931) on $Ki^{ref}$ in the BMI subsample. The figure shows posterior distributions with medians (point) and intervals (95% thin and 80% thick line) on a standardized scale.

The PD effect on thalamic $Ki^{ref}$ was positive only in the model using the total sample and not analyzing the BMI effect, while the effect was negative in the BMI subsample whether the BMI effect was estimated or not. Thus, the change in the PD effect must stem from the sample cut, rather than adding BMI as a predictor. This was supported by the assessment of the thalamic $Ki^{ref}$ estimates in the remaining subjects (n= 130, 75 males and 55 females) who were cut from the BMI subsample (**Figure S6**). While the groups are not clearly separable by $Ki^{ref}$, there are a few healthy controls with notably low $Ki^{ref}$ estimate, likely causing the positive PD effect in the model using the total data. Despite that most of the young subjects are healthy controls, the age and sex distributions of the subject groups appear rather balanced.

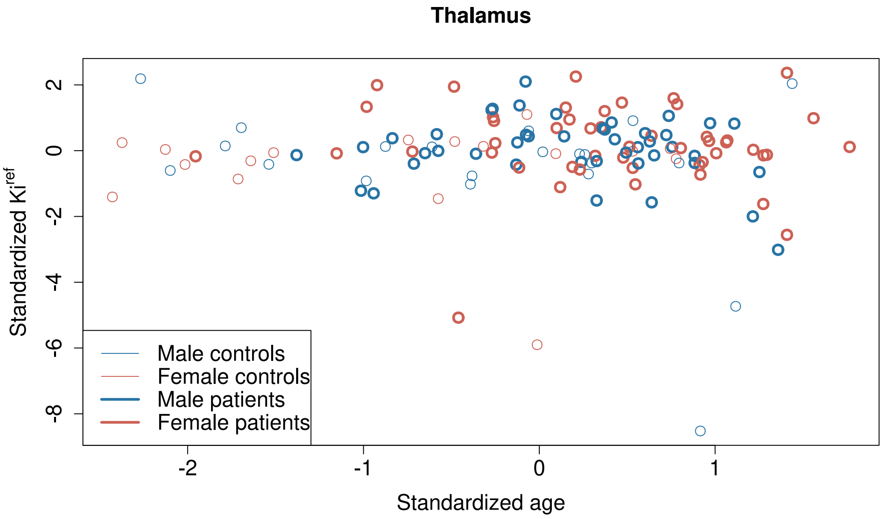


**Figure S6**. Standardized thalamic $Ki^{ref}$ estimates in the subjects excluded from the BMI subsample (n= 130, 75 males and 55 females). The observations are shown across standardized age in thalamus, separately for the groups and sexes. $Ki^{ref}$ and age are standardized to the total sample (0= mean and 1= SD in the total sample) to remain their comparativeness to the total sample.

### Model diagnostics

*The effect of priors on the regression model findings*

The effect of priors on model findings decrease with increasing sample size (McElreath, 2020), while we had a large sample of hundreds. However, we wanted to assess how sensitive our regression modeling was to our priors by testing alternative priors.

Based on previous literature, dopamine synthesis capacity is lowered in high BMI (Janssen & Horstmann, 2022), in males versus females (Laakso et al., 2002), and in PD patients versus healthy controls (e.g. Cropley et al., 2008; Pavese et al., 2011; Samii et al., 1999). Due to the lack of independent studies investigating the sex difference in the dopamine synthesis capacity, some of our data may overlap with the data used in the work by Laakso and colleagues (Laakso et al., 2002), although the priors and data should be independent sources of information (McElreath, 2020). As a result, the prior may be partially dependent on the data analyzed here. However, the data used in the work by Laakso and colleagues (Laakso et al., 2002) was only 35 subjects (23 males and 12 females), making it at maximum (if all the subjects were included here) only about 16% of the BMI subsample used in the analysis. As our alternative priors, we accordingly set normal distribution with expected value of -1 and standard deviation (SD) of 0.5 to the effects of standardized BMI (slope), sex (males versus females), and PD (PD patients versus healthy controls).

In the alternative prior, the expected value is negative, assigning more probability on the negative than positive regression coefficients, unlike the primary prior with the expected value of 0, assigning equal probabilities to positive and negative regression coefficients. However, as normal distributions, both are continuous excluding no regression coefficients. The alternative prior is narrower (SD = 0.5) than the primary one (SD = 1). When data is normally distributed, around 95% of the observations are within |2 * SD| from the mean and median (Nummenmaa, 2021). This means that with the prior SD of 0.5, around 95% of the alternative prior probability is assigned to the regression coefficients from -2 to 0 (compare from -2 to 2 in the primary prior). The remaining predictors standardized age, and scanner were given the same normal prior (expected value = 0, SD = 1) as in the primary modeling. The primary and the alternative priors are visualized in **Figure S7**.

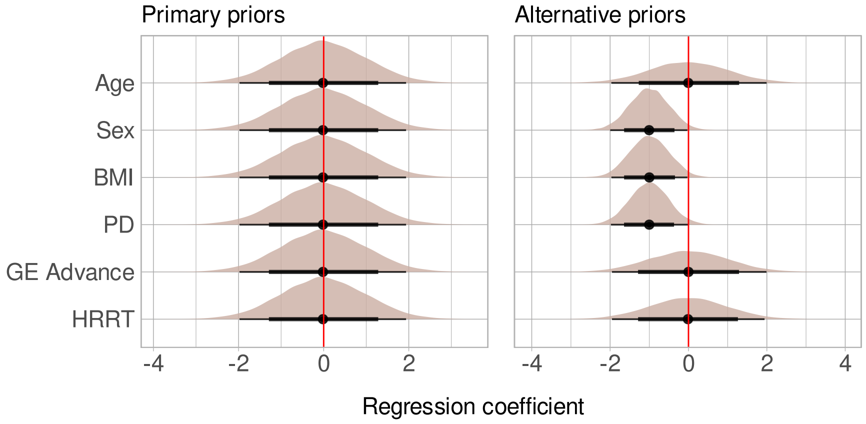


**Figure S7**. Primary (left) and alternative (right) prior distributions, assigned to the main effects of age, sex (males versus females), BMI, PD (PD patients versus healthy controls), and scanner (each scanner versus Ecat 931) on $Ki^{ref}$. The figure shows the prior distributions with medians (point) and intervals (95% thin and 80% thick line) on a standardized scale.

We did not alter any other model specifications than the priors. As data, we used the subsample of subjects with the BMI information available (BMI subsample, n= 220), as the BMI effect was included in the modeling. The models showed no divergent transitions nor Rhats > 1, supporting sufficient model convergence (Vehtari et al., 2021). The findings of the regional models are given in **Figure S8**. The findings of the corresponding models using the primary and the alternative priors are essentially the same. As a conclusion, our primary findings are not highly sensitive to the primary priors when compared with the alternative priors described here.

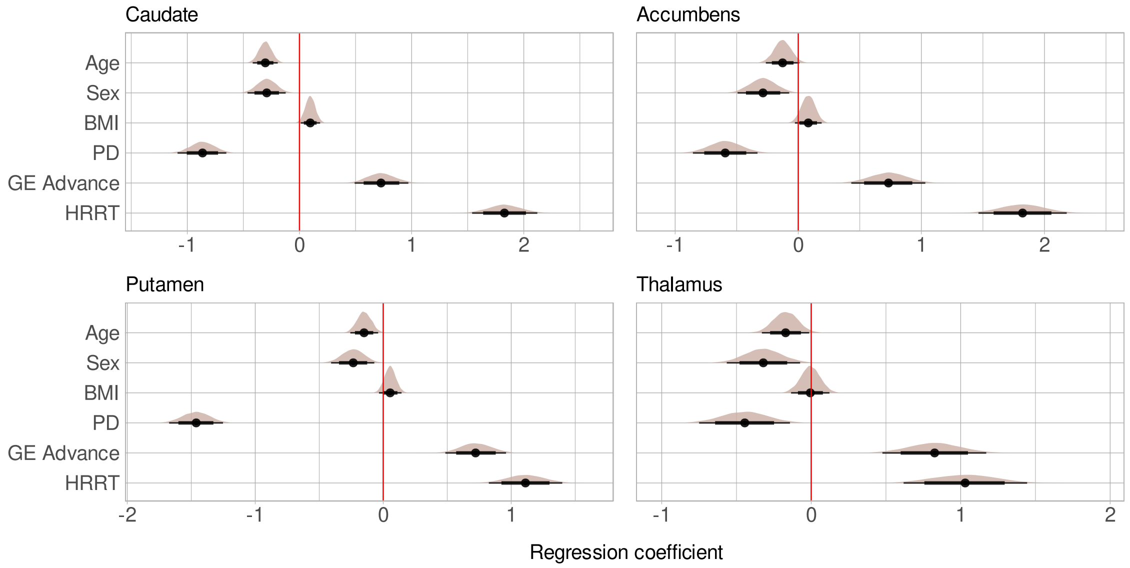


**Figure S8**. Main effects of age, sex (males versus females), BMI, PD (PD patients versus healthy controls), and scanner (each scanner versus Ecat 931) on $Ki^{ref}$ in the BMI subsample, using the alternative priors. The figure shows posterior distributions with medians (point) and intervals (95% thin and 80% thick line) on a standardized scale.

*Residuals*

As model diagnostics, we assessed the normality and the homoscedasticity of the residuals by visualizing the fitted values on the residuals from each regional model including standardized age, sex, standardized BMI, PD, and scanner as predictors and restricting the relationships between each predictor and the dependent variable (standardized $Ki^{ref}$) as linear (**Figure S9**).

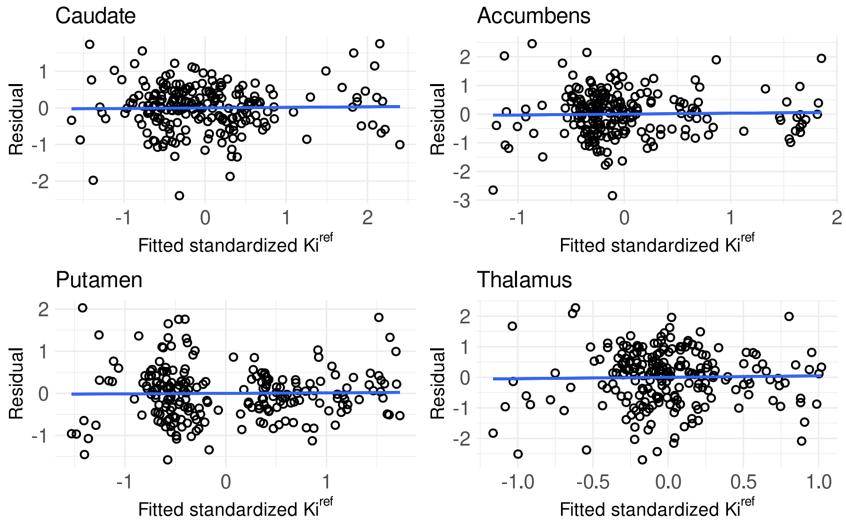


**Figure S9**. Fitted standardized $Ki^{ref}$ (x-axis) on the residual (y-axis) in each regional model including standardized age, sex, standardized BMI, PD, and scanner as predictors and restricting the relationships between each predictor and the dependent variable (standardized $Ki^{ref}$) as linear. The blue line reflects the fit, i.e. where residual = 0.

None of the regional models show clear signs of heteroscedasticity, as the residual sizes appear rather stable across the fitted values. The residuals also appear randomly distributed along the fit (residual= 0), and no clear systematic structures of the residuals are spotted, further supporting sufficient model fit.

### Interaction of age and sex

To make sure we did not overlook possible sex differences in the age effects, we ran regional models where the age effect was estimated separately in males and females (interaction of age and sex). The models were otherwise identical to the models where all predictors (including BMI) were involved. Overall, the age effect was negative in both males and females (**Figure S10**). In accumbens, however, the age-effect was more prominent in females than males.

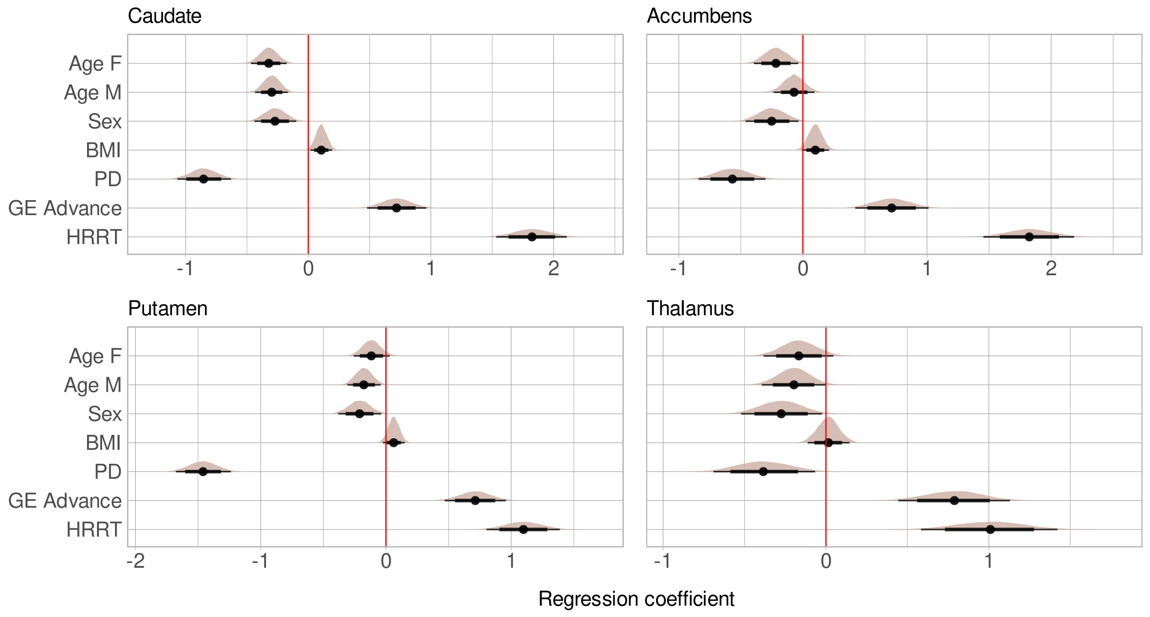


**Figure S10**. Interaction of age and sex (males versus females), and main effects of BMI, PD (PD patients versus healthy controls), and scanner (each scanner versus Ecat 931) on $Ki^{ref}$ in the subsample of subjects with BMI information available (BMI subsample, n= 220), using alternative priors. Age F (M)= Age effect in females (males). Sex= the effect of sex (males versus females) at sample mean age (55 years on original scale, 0 on standardized scale). The figure shows posterior distributions with medians (point) and intervals (95% thin and 80% thick line) on a standardized scale.

### Linearity assessment of the age and BMI effects

To make sure we did not underfit the effects of age and BMI on $Ki^{ref}$ (all standardized) by approximating them with a linear function, we ran additional regional regression models allowing nonlinearity in these effects. Based on the documentation (https://paulbuerkner.com/brms/reference/s.html), we used smooth terms (s) within model formulas (Wood, 2004). To assess the possible non-linearity of both age and BMI effects in the same region-specific models, we used the BMI subsample.

To facilitate the converge of these models that are more complex than the primary ones, we set adapt-delta to 0.999 and maximum treedepth to 20, based on instructions in https://mcstan.org/misc/warnings.html#divergent-transitions-after-warmup. We used the primary priors described in the manuscript. With these specifications, the models were free from divergent transitions, and Rhats > 1, supporting sufficient convergence of the models (Vehtari et al., 2021).

The estimated effects allowing nonlinearity (splines) are illustrated in **Figure S11**. Particularly in caudate and accumbens, the findings suggest that in early adulthood, there may be a positive or nonexistent age – $Ki^{ref}$ association that becomes negative approximately at the age of 50. The uncertainty in the age effects appears greater in the early versus late adulthood (posterior intervals wider at younger versus older ages), probably reflecting the number of observations throughout the age range (stronger representation of older versus younger subjects). Despite the wide posterior intervals indicating uncertainty, particularly the striatal ROIs show subtle support for positive relationship between BMI and $Ki^{ref}$, and there is no clear support for nonlinearity in these effects. The remaining effects remained roughly the same in the models that allowed nonlinearity in the effects of age and BMI (**Figure S12**).

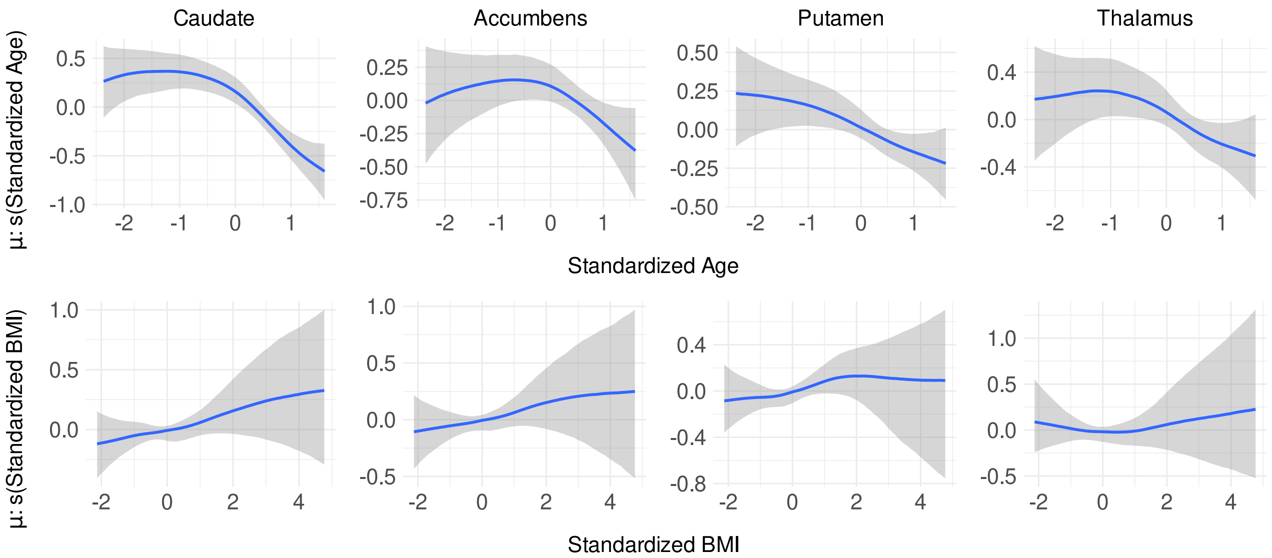


**Figure S11**. The effects of age (upper panel) and BMI (lower panel) on $Ki^{ref}$ as splines in the subsample of subjects with BMI information available (BMI subsample, n= 220). The figure shows the partial effects of the predictors (x-axis) on the expected $Ki^{ref}$ (μ: s(predictor) on the y-axis) as posterior medians (blue spline) and 95% intervals (shaded areas) on a standardized scale. In the upper panel, x= 0 is the mean age= 55 years, and one unit is one standard deviation (SD)= 15 years in the sample. Similarly, in the lower panel, x= 0 is the mean BMI= 26, and one unit is one SD= 4 in the sample.

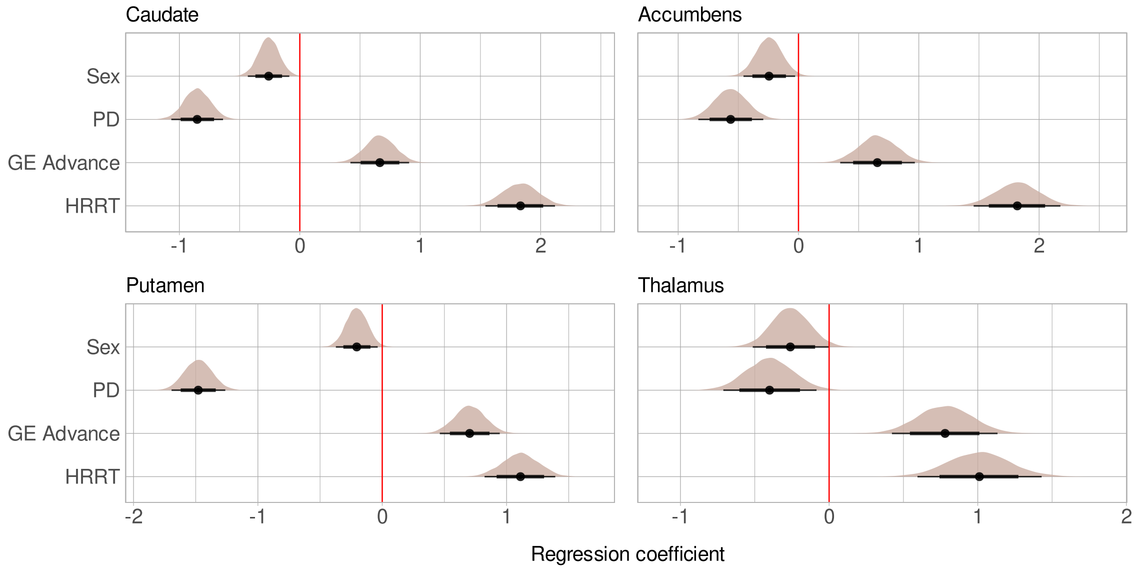


**Figure S12**. Main effects of sex (males versus females), PD (PD patients versus healthy controls), and scanner (each scanner versus Ecat 931) on $Ki^{ref}$ in the BMI subsample in the models allowing nonlinearity in the age and BMI effects. The figure shows posterior distributions with medians (point) and intervals (95% thin and 80% thick line) on a standardized scale.
